# Arousal state alters brain network switching and moderates cognitive task performance

**DOI:** 10.64898/2026.03.09.710652

**Authors:** Kimberly Kundert-Obando, Haatef Pourmotabbed, Kamalpreet Kaur, Shiyu Wang, Juan Gomez Lagandara, Sarah E. Goodale, Caroline Martin, Victoria L. Morgan, Dario J. Englot, Lucina Q. Uddin, Mikail Rubinov, Catie Chang

## Abstract

Arousal relates to cognitive performance, but the neural underpinnings of this relationship remain unclear. One candidate marker is switching rate, a dynamic measure that has been linked to cognition and has been speculated to be sensitive to arousal. However, whether switching rate is altered across arousal states has not been directly tested. Here, using fMRI together with concurrent eye monitoring and EEG, we examined how the switching rates of the default mode, salience, and central executive networks are altered across arousal states. Default mode and anterior salience networks exhibited significant differences in switching rates across arousal states determined with eye tracking. Notably, thalamic subregions showed arousal-dependent changes in switching rate that were replicated across independent datasets and arousal measures. Additionally, arousal moderated the relationship between average network switching and performance on a relational processing task. Together, these findings suggest that switching rate may index neural underpinnings of arousal-dependent cognition.

## Introduction

Arousal refers to a spectrum of states varying between alert wakefulness, drowsiness, and sleep (Jones, 2003). Levels of arousal constantly fluctuate throughout the course of a day, and are thought to influence cognitive task performance (Alameda et al., 2024; Barber et al., 2020; Estarellas et al., 2024). The Yerkes-Dodson law states that arousal is related to cognition in an inverted-U manner, where either extremely high (hyper-arousal) or low (hypo-arousal) levels of arousal impair cognition, while a median level of arousal supports optimal cognitive performance (Yerkes & Dodson, 1908). Indeed, many studies support this law, or reveal a more complex relationship between arousal and cognition, in domains such as working memory (Hansen et al., 2003; Heitz et al., 2008; Robison et al., 2023; Robison & Brewer, 2020; Unsworth & Robison, 2017), cognitive control (Alameda et al., 2024; van der Wel & van Steenbergen, 2018; Zhang et al., 2012), and attention (Barber et al., 2020; Foucher et al., 2004). Further, disruption of the arousal system may underlie cognitive impairment in early-stage dementia (Estarellas et al., 2024; Jacobs et al., 2015), schizophrenia, autism, and depression (Mathersul et al., 2013; Suttkus et al., 2021; Xie et al., 2024). Therefore, investigating the neural underpinnings behind arousal-dependent cognitive changes is crucial for uncovering potential biomarkers of cognitive impairment across neurodegenerative and psychiatric disorders.

Prior work provides evidence that network switching is a brain marker of cognition. Network switching, also known as network flexibility, is a measure that quantifies the dynamic interactions between brain nodes across time. When two or more nodes are strongly connected, they form a community – much like how people who frequently interact form social communities. However, neural community assignments can switch across time, and the overall amount by which a given node switches its community is termed network switching (Figure 1). In functional magnetic resonance imaging (fMRI) studies, network switching has been linked with multiple cognitive functions, including attention, learning, emotion, working memory, and optimal cognitive performance (Bassett et al., 2011a; Braun et al., 2015a; Pedersen et al., 2018; Shine, Bissett, et al., 2016a). Interestingly, fatigue and sleep duration have been reported to influence network switching (Betzel et al., 2017; Pedersen et al., 2018; Shine, Bissett, et al., 2016a), leading to the view that arousal state may be a confound in prior studies relating network switching to cognition.

**Figure 1.**
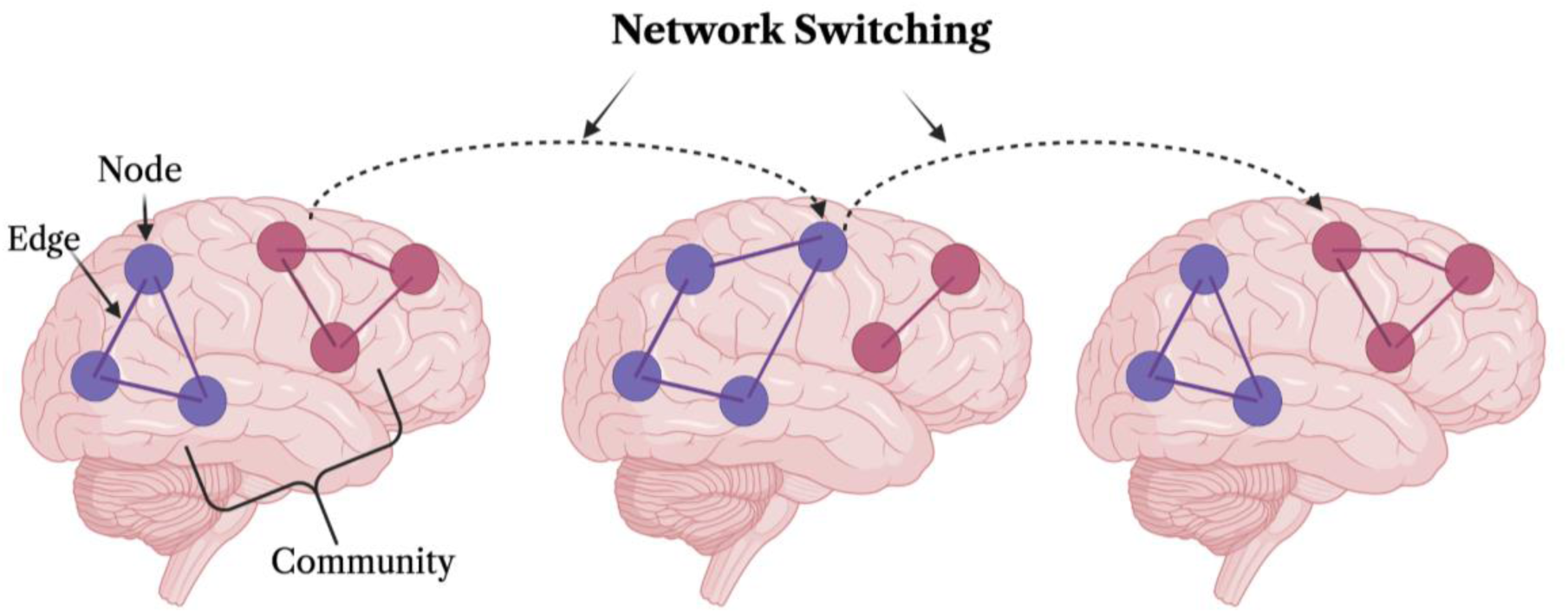
Illustration of Network Switching. Network Switching is defined as the number of times per scan in which a given node switches its community assignment. Each circle represents a node (which can comprise either an entire network or a single parcel) and the color corresponds to the community assignment. The switching of a node between communities is indicated with the dashed arrow. Figure generated using BioRender.com

However, we propose a different perspective: that the coupling between arousal and cognition may position network switching as a potential brain marker of arousal-dependent cognitive changes. To test this hypothesis, a first critical step is to determine if network switching varies with arousal in the absence of a task. As prior studies that related network switching and arousal during rest have relied on self-reported fatigue or sleep duration measures (Betzel et al., 2017; Pedersen et al., 2018), there is a need to investigate whether network switching rate relates to concurrent objective measures of arousal. Moreover, an important consideration is that several basic characteristics of the fMRI signal are known to be sensitive to changes in arousal state. For example, drowsiness is accompanied by increased signal fluctuation amplitude and spatial correlations across the brain, along with altered cross-network correlations (Chang et al., 2013; Fukunaga et al., 2006; Haimovici et al., 2017; Liu & Falahpour, 2020). Therefore, it is necessary to investigate the extent to which network switching varies with arousal beyond these more basic fMRI characteristics. Lastly, to assess whether network switching indexes arousal-dependent cognition, it is important to determine whether arousal state moderates established relationships between network switching and cognitive performance (Bassett et al., 2011; Betzel et al., 2017; Braun et al., 2015; Pedersen et al., 2018).

To address these gaps, here we investigate whether the switching dynamics of three major brain networks—the default mode, salience, and central executive networks (Menon, 2011; Uddin, 2015)—vary with concurrent measures of arousal (electroencephalography [EEG] and eye monitoring) at rest in healthy adults. We focus on these networks because they exhibit broad, coordinated engagement across a range of cognitive tasks (Chand et al., 2017; Oyarzabal et al., 2022; Suttkus et al., 2021; Unsworth & Robison, 2017) and their coordination may be influenced by arousal (Young et al., 2017). In addition, we adopt integrative statistical modeling to test whether network switching indexes arousal more sensitively than several basic characteristics of the fMRI signal (Rubinov, 2025; Váša & Mišić, 2022). Further, motivated by theories that arousal initiates the salience network to coordinate the activation and deactivation of the central executive and default mode networks, respectively (Unsworth & Robison, 2017), we also investigate if the amount of time in which the salience network shares its community with the central executive or default mode network is altered across arousal states. Lastly, we tested if arousal state impacts how global (average) brain network switching, and other arousal-sensitive fMRI characteristics, relates to cognitive performance.

## Results

This study drew upon simultaneous EEG-fMRI data collected at Vanderbilt University (“VU-EEG-fMRI”), and simultaneous eye tracking-fMRI data collected in the Human Connectome Project 7T dataset (“HCP-7T”). Both EEG (Lendner et al., 2020) and eye tracking (Gonzalez-Castillo et al., 2022) provide validated measures of arousal, which motivated our selection of these datasets. Our investigation focused on the default-network (DMN), salience network (SAL), and central executive network (CEN), which were delineated based on a published functional network atlas (FINDLAB Atlas; Shirer et al., 2011). This atlas further partitions the DMN into dorsal and ventral subnetworks (DDMN, VDMN), the SAL into anterior and posterior subnetworks (ASAL, PSAL), and the central executive network into left and right subnetworks (LCEN, RCEN).

### Network switching across arousal state

First, we investigated whether the switching rates of these networks of interest changed across arousal states. Our hypothesis regarding the salience network centers on its role in detecting salient stimuli and coordinating the activity of large-scale networks (Menon & Uddin, 2010; Uddin, 2016). Given this role, together with the arousal-biased competition theory – which posits that higher arousal increases the priority of salient stimuli – we hypothesized that higher arousal would lead to higher switching of the salience network dynamic activity (Kim et al., 2024). In contrast, the activation of the default mode network may result from a reduction of cholinergic release (Sanda et al., 2024); therefore, we may expect its switching to be elevated in lower states of arousal (drowsy states), where cholinergic release is typically lower (Teles-Grilo Ruivo et al., 2017). Since the switching of the central executive network is strongly associated with cognition during a task (Braun et al., 2015a), we hypothesized that this network would show an arousal dependence only during a task and not at rest.

In the HCP-7T data, after FDR corrections (q=0.05) across networks of interest, the anterior salience network showed a significant increase of switching in alert states (z=-3.23, r=-0.17, q=0.004). Both the dorsal and ventral component of the default mode network showed increased switching in drowsy states (DDMN: z=3.20, r= 0.17, q=0.004; VDMN: z=2.49, r=0.13, q=0.03; **Figure 2**). These results did not reach statistical significance in the VU-EEG-fMRI dataset after FDR corrections (q=0.05). All network-level statistics are reported in **Supplementary Table 1a**. When replicating the investigation at the parcel level, statistical analysis of the HCP-7T dataset indicated that thalamic parcels within the default mode and central executive networks had significantly higher switching rates in alert states (DDMN bilateral thalamus: z=-6.25, r=-0.34, q<0.001; and RCEN thalamus-caudate: z=-7.08, r=-0.38, q<0.001). These two findings were replicated in the VU-EEG-fMRI data (DDMN bilateral thalamus: z=-3.53, r=0.75, q=0.02; RCEN thalamus-caudate: z=-3.19, r=-0.68, q=0.03). Additionally, several parcels across all networks of interest (DDMN, VDMN, RCEN, LCEN, ASAL, and PSAL) had significantly different levels of switching across arousal states in the HCP-7T data only (**Figure 3**). For all parcel-level statistics, please refer to **Supplementary Table 1b**.

**Figure 2.**
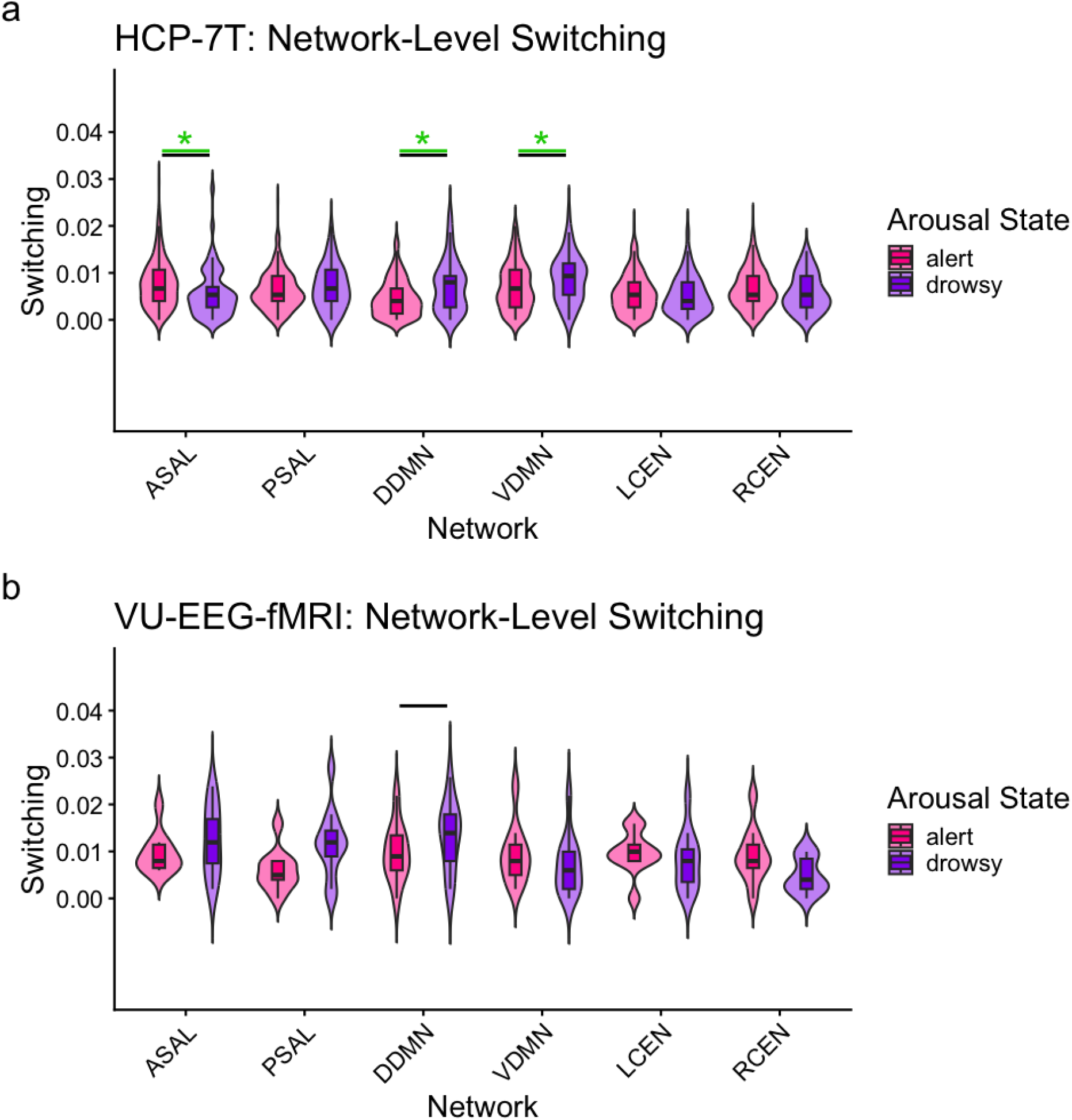
Network switching in alert and drowsy arousal states. a) Network-level switching across arousal states in the HCP-7T data. b) Similar plots for VU-EEG-fMRI data. Violin plots show switching values for each large-scale brain network during alert (pink) and drowsy (purple) states. Green bar represents significance after Mann–Whitney U-test (where * indicates p < 0.05), and a black bar indicates that the finding was significant through the third null model. Network abbreviations: Anterior Salience Network (ASAL), Posterior Salience Network (PSAL), Dorsal Default Mode Network (DDMN), Ventral Default Mode Network (VDMN), Left Central Executive Network (LCEN), Right Central Executive Network (RCEN).

**Figure 3.**
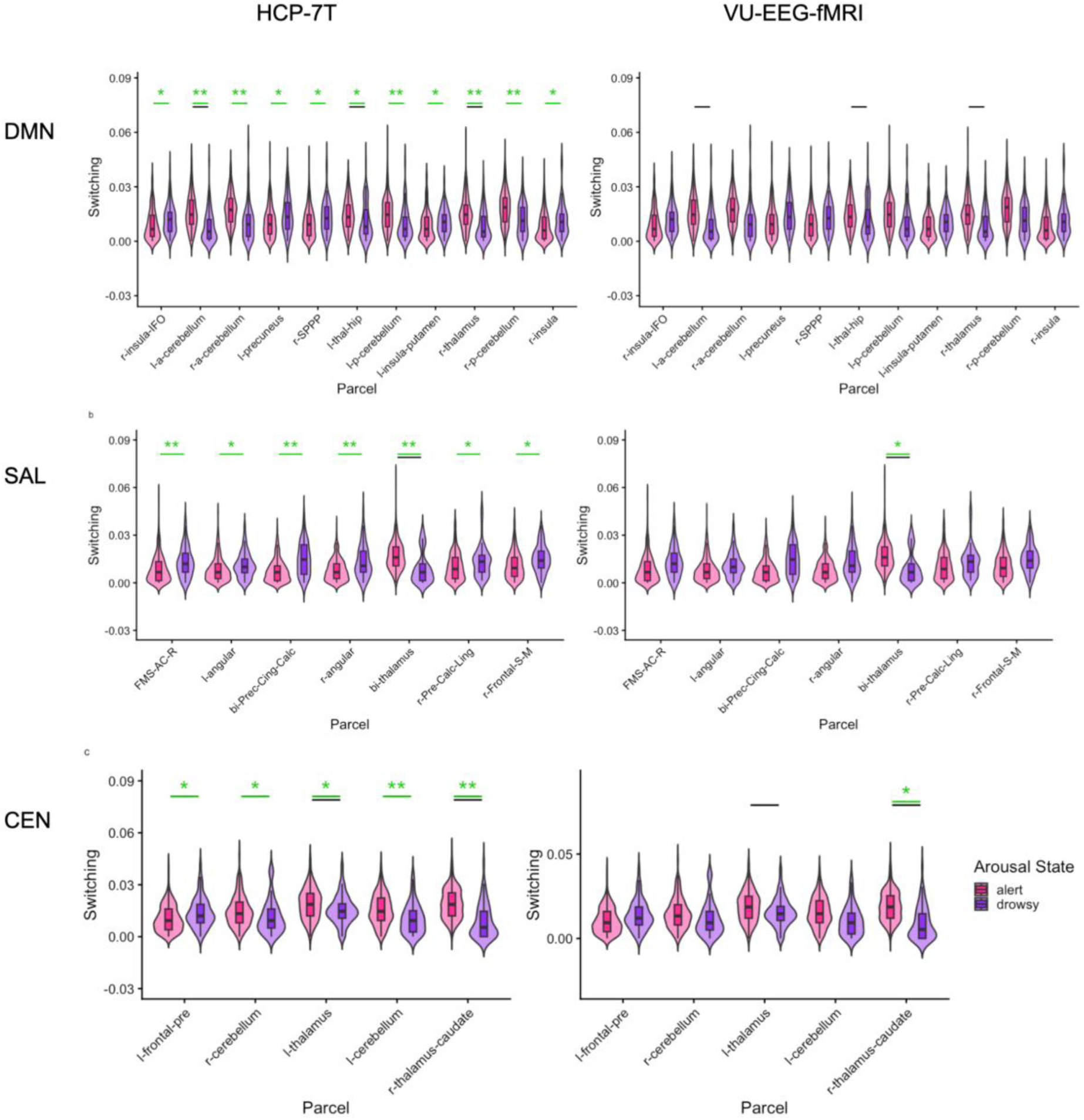
Parcel-level switching in alert and drowsy arousal states. a–e) Parcel-level switching across arousal states in the HCP-7T (left) and VU EEG-fMRI dataset (right). a-b DDMN, VDMN). c-d (PSAL-ASAL), e-f (LCEN, RCEN). Violin plots show parcel-level flexibility across two arousal states: alert (pink) and drowsy (purple). Significant differences between arousal states are indicated with a green bar and with asterisks for *p < 0.05, **p < 0.001 (Mann-Whitney U-test). A black bar indicates that the finding was significant through the third null model. The following acronyms are used for parcel names: r-insula-IFO (right insula/inferior frontal operculum), l-a-cerebellum (left anterior cerebellum), r-a-cerebellum (right anterior cerebellum), l-precuneus (left precuneus), r-SPPP (right superior posterior parietal cortex), l-thal-hip (left thalamus/hippocampus), l-p-cerebellum (left posterior cerebellum), l-insula-putamen (left insula/putamen), r-thalamus (right thalamus), r-p-cerebellum (right posterior cerebellum), r-insula (right insula), FMS-AC-R (frontal medial superior/anterior cingulate, right), l-angular (left angular gyrus), bi-Prec-Cing-Calc (bilateral precuneus/cingulate/calcarine cortex), r-angular (right angular gyrus), bi-thalamus (bilateral thalamus), r-Pre-Calc-Ling (right precuneus/calcarine/lingual gyrus), r-Frontal-S-M (right superior medial frontal gyrus), l-frontal-pre (left prefrontal cortex), r-cerebellum (right cerebellum), l-thalamus (left thalamus), l-cerebellum (left cerebellum), and r-thalamus-caudate (right thalamus/caudate)

Next, we tested the extent to which arousal-dependent changes in network switching persist after controlling for changes in several basic fMRI signal characteristics (**Supplementary Table 2a**). To this end, we designed four null models that progressively preserved the following fMRI signal characteristics: (1) mean across time and across nodes, (2) variation across time and across nodes, (3) correlation between each node and the global mean signal, and lastly, (4) static correlation between nodes of interest (Gonzalez et al., 2019; Rubinov, 2016, 2023, 2025; Váša & Mišić, 2022). In the HCP-7T data, and mirroring the results of the above statistical tests, arousal-dependent switching of default mode subnetworks (DDMN, p<0.001; VDMN, p<0.001), and the anterior salience network (ASAL, p<0.001) persisted after controlling for spatial and temporal mean and variance as well as correlation with the global signal (null models 1-3). However, these effects were no longer significant when additionally maintaining the static correlation between salience, default mode, and central executive networks (null model 4). In other words, arousal-dependent changes in the switching of all three networks could arise from arousal-dependent changes in the strength of their static correlation. In the VU-EEG-fMRI data, DDMN survived the tests against null models 1-3, but did not survive when static correlation was preserved (null model 4). For the parcel-level investigation, thalamic parcels associated with multiple networks survived null models 1-3 – specifically, those associated with PSAL (p=<0.001), DDMN (p<0.001), and LCEN (p<0.001) (**Figure 3**). Interestingly, the RCEN caudate-thalamus parcel survived up to null model 4 (p<0.001) in both datasets, suggesting that the switching behavior of this parcel carries unique, arousal-related information not explained by the fMRI signal characteristics encoded in the null models (**Supplementary Table 2b**).

### Static correlation and network-to-global-mean correlation across arousal state

As the null models revealed, static correlation may be a key factor influencing the arousal dependency of network switching. Moreover, prior work has reported a strong arousal dependence in the magnitude and spatial distribution of the global mean signal (Liu & Falahpour, 2020). Therefore, we next sought to examine the behavior of the mean static correlation (the average pairwise correlation across the 14 sub-networks), as well as the correlations between ASAL, DDMN, and VDMN with the global mean signal, across arousal states. Static correlation was significantly altered across arousal states in the HCP-7T dataset but not in the VU-EEG-fMRI dataset (z=4.32, r=0.23, q<0.001; z=1.71, r=0.37, q=0.08). For correlations with the global mean signal, DDMN was significant in both HCP-7T and VU-EEG-fMRI datasets (z=4.80, r=0.26, q<0.001 and z=2.77, r=0.59, q=0.02); VDMN was significant in HCP-7T but not in VU-EEG-fMRI (z=7.06, r=0.38, q<0.001 and z=1.98, r=0.42, q=0.06); and ASAL was significant in VU-EEG-fMRI but not in HCP-7T (z=2.44, r=0.52, q=0.03 and z=-1.16, r=-0.06, q=0.23). Statistical details of these findings are reported in **Table 1**.

**Table 1.** fMRI connectivity measures across arousal states. Mann-Whitney U test statistics for testing arousal-state differences in average static correlation (STAT-COR) and for testing arousal-state differences in the correlation of the default mode network (DMN), ventral default mode network (VDMN), and anterior salience networks (ASAL) with the global signal (GS).

| HCP-7T |  |  |  |  |  |  |  |
| --- | --- | --- | --- | --- | --- | --- | --- |
| Network | U | RankMean<br>Drowsy | RankMean<br>Alert | *z value* | *r value* | *p value* | *q value* |
| DDMN-GS | 11963.00 | 229.88 | 161.67 | 4.80 | 0.26 | <0.001 | <0.001 |
| VDMN-GS | 13553.00 | 256.38 | 156.11 | 7.06 | 0.38 | <0.001 | <0.001 |
| ASAL-GS | 7765.00 | 159.92 | 176.35 | -1.16 | -0.06 | 0.25 | 0.25 |
| STAT_COR | 11623.00 | 224.22 | 162.86 | 4.32 | 0.23 | <0.001 | <0.001 |
| VU-EEG-fMRI |  |  |  |  |  |  |  |
| Network | U | RankMean<br>Drowsy | RankMean<br>Alert | *z value* | *r value* | *p value* | *q value* |
| DDMN-GS | 102.00 | 15.00 | 7.30 | 2.77 | 0.59 | 0.01 | 0.02 |
| VDMN-GS | 90.00 | 14.00 | 8.50 | 1.98 | 0.42 | 0.05 | 0.06 |
| ASAL-GS | 97.00 | 14.58 | 7.80 | 2.44 | 0.52 | 0.01 | 0.03 |
| STAT_COR | 86.00 | 13.67 | 8.90 | 1.71 | 0.37 | 0.09 | 0.09 |

### Salience Network Community Allegiance across Arousal State

Menon and Uddin proposed that the salience network plays a role in engaging the central executive network, and disengaging the default mode network, to facilitate goal-directed cognitive functions (Menon & Uddin, 2010; Uddin, 2015). Later work extended this theory by suggesting that arousal circuitry – specifically, locus coeruleus noradrenergic projections – enhances neuronal activity to detect salient stimuli, facilitating the function of the salience network (Unsworth & Robison, 2017). Motivated by both theories, we asked if arousal state alters the fraction of time in which the salience network shares a community with either central executive network or default mode network (i.e., “shared community allegiance”). We hypothesized that salience and central executive networks would exhibit shared allegiance more often during higher levels of arousal, as it is thought that sustained arousal increases salience and central executive network interactions to promote optimal cognition (Unsworth & Robison, 2017). In contrast, we suspected that the allegiance between salience and default mode networks would emerge as stronger during drowsy states, based on prior observations that the coupling between key nodes of these networks (posterior cingulate cortex, anterior cingulate cortex) was elevated during rest compared to during a visual processing task (Greicius et al., 2003).

In the HCP-7T dataset, all significant pairs of SAL-DMN subnetworks, rank-based probabilities were higher in drowsy compared to alert states, supporting our hypothesis. (e.g., ASAL–DDMN: z=3.46, r=0.18, q<0.001; PSAL–DDMN: z=3.88, r=0.21, q<0.001; PSAL-VDMN: z=2.11, r=0.11, q=0.04). These results were replicated in the VU-EEG-fMRI data. Contrary to our hypothesis, subnetworks of the salience and central executive networks in the HCP-7T data were found to have significantly higher shared allegiance in the drowsy, compared to alert state (ASAL-LCEN: z=3.191, r=0.17, q<0.001; ASAL-RCEN: z=4.75, r=0.26, q<0.001; PSAL-LCEN: z=2.03, r=0.11, q=0.04; PSAL-RCEN: z=3.06, r=0.16, q=0.003). This did not replicate in the VU-EEG-fMRI data **(Figure 4)**.

**Figure 4.**
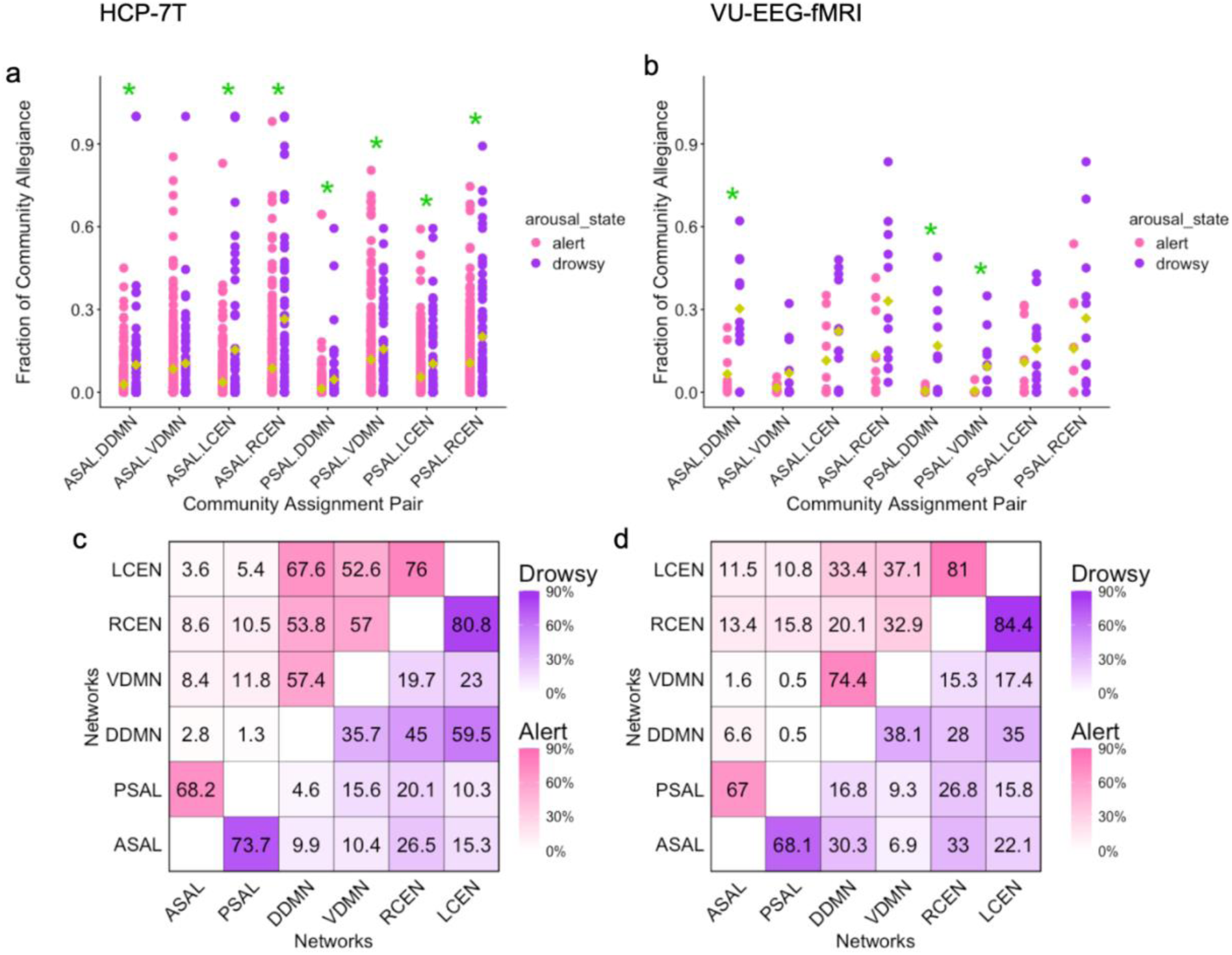
Arousal dependence of network community allegiance. a, b) The fraction of time in which networks X and Y share a community assignment (allegiance) is denoted as X.Y, and is shown for a) the HCP-7T data and b) the VU-EEG-fMRI data. Green asterisk indicates that community allegiance was significantly different across arousal state q<0.05. c, d) Heat map indicates the percentage of time windows for which two networks share a community (allegiance), for drowsy (purple) scans and alert (pink) scans, shown for the c) HCP-7T data and d) VU-EEG-fMRI data. Networks investigated were as follows: posterior salience network (PSAL), anterior salience network (ASAL), left central executive network (LCEN), right central executive network (RCEN), dorsal default mode network (DDMN), and ventral default mode network (VDMN).

### Global brain switching across arousal state

Global (“whole-brain”) switching, defined as the average switching across nodes at a network level, has been found to have clinical relevance as it is consistently reported to be higher in schizophrenia (Braun et al., 2016; Gao 2024; Wei 2022). To understand if previous findings may have been influenced by arousal state, here we tested whether global brain switching is sensitive to arousal state. We found no significant effects in either the HCP-7T (U=8195.5, z=-0.55, r=-0.03, p=0.58) data or VU-EEG-fMRI dataset (U=53, z=-0.46, r=-0.10, p=0.67).

### Arousal state moderates how global brain switching relates to task accuracy

Alongside its potential clinical relevance, global brain switching has been found to predict cognitive performance. Specifically, prior work (Pedersen et al., 2018) demonstrated that global brain network switching predicted working memory task performance and relational task performance in the HCP-3T dataset. However, while the HCP-3T dataset contains these task performance measures, it does not have validated arousal measures. Since eye monitoring is included with the HCP-7T resting-state data, we use the HCP-7T data (acquired in the same subjects) to investigate if the relationship between global brain switching and task performance is moderated by arousal state. After FDR correction for multiple comparisons, we found that arousal state moderated the relationship between global network switching and relational task accuracy (β = –5090.17, SE=1889.7, q=0.01), but did not moderate its relationship with working memory task accuracy (β = –1257, SE= 1132.7, q=0.27); **Figure 5(a,b)**. Specifically, for individuals in an alert state during the scan when global network switching was measured, higher performance was related to higher global switching rate; whereas for those in the drowsy state when switching was measured, worse performance was related to higher switching. These findings provide preliminary evidence that global brain switching may be a marker of certain arousal-dependent cognitive changes.

**Figure 5.**
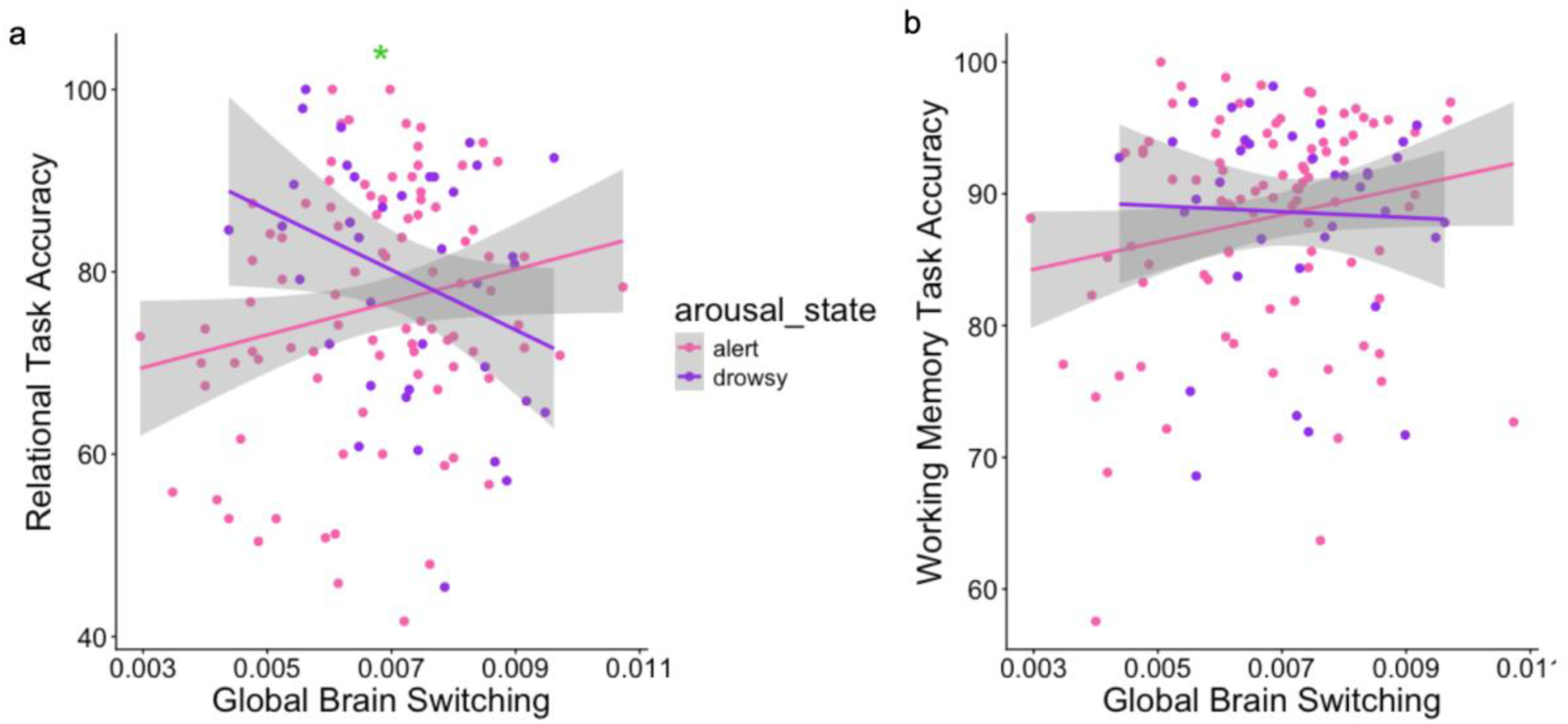
Arousal state moderates the association between global network switching and relational task performance. a) Arousal state was found to moderate the relationship between global brain switching and relational task accuracy (β = –5090.17, q=0.01). b) Arousal state did not significantly moderate the relationship between global switching and working memory task accuracy (β = –49.5, q=0.2)

### Arousal state does not moderate how static correlation relates to task accuracy

Since the null model analysis revealed a prominent contribution of static correlation to arousal-dependent network switching, we further tested whether arousal moderates the relationship between static correlation and task accuracy. We found that arousal did not significantly moderate the relationship between static correlation and relational task performance (β = –115.3, SE = 59.2, q = 0.28) or working memory task accuracy (β = –49.5, SE = 36.3, q = 0.28).

## Discussion

Transitions of arousal state are accompanied by altered cognition; however, brain markers of arousal-dependent cognitive changes have not been fully characterized. Network switching is an established marker of cognition and has been speculated to be sensitive to arousal state (Bassett et al., 2011a; Pedersen et al., 2018; Shine, Bissett, et al., 2016a), though the latter has not been directly tested with objective measures of arousal. The present study sought to determine if the switching dynamics of three networks – default mode, salience, and central executive networks – is altered across arousal state. We used fMRI data, recorded concurrently with EEG and eye tracked-based arousal measures, to demonstrate that the switching dynamics of the default mode and salience networks – and of thalamic subregions – is sensitive to arousal state. Moreover, we used integrative modeling to identify which fMRI signal characteristics promote these arousal-state dependencies. We also show that arousal state impacts how frequently the salience network shares a community with default mode and central executive networks. Lastly, we show that arousal state moderates the relationship between global brain switching and performance on a relational processing task. Overall, our findings indicate that network switching may index arousal-dependent cognitive changes.

We first sought to determine if the network switching of the salience, default mode, and central executive networks is altered across arousal states. Consistent with our hypothesis, the salience network showed an increase in its spontaneous switching rate in alert states. This finding may arise from how the salience network interacts with the locus coeruleus, a key brain region that regulates arousal levels. We speculate that alert states may promote phasic firing of the locus coeruleus, which is known to increase during moments of salience, thereby facilitating the switching of the salience network (Aston-Jones & Cohen, 2005; Grimm et al., 2024). Our findings also provide support for the theories proposed by Unsworth and Robinson (2017), which suggest that arousal modulates the salience network’s interactions with other large-scale brain networks. In terms of the default mode network, we hypothesized that this network would show greater switching in drowsy states in rest, given evidence that cholinergic activity—elevated during drowsiness—promotes its activation (Kim et al., 2024). Indeed, we observed that the default mode increases its dynamic connections with other networks at lower arousal levels, which may be inherent to its role in facilitating internal processing, such as mind wandering (Bassett et al., 2011a; Fransson, 2005). Lastly, we suspected that any arousal-dependent switching of the central executive network would only emerge if a task was presented to participants. It has been previously established that the switching of the central executive network is elevated during task engagement and stable at rest (Braun et al., 2015b; Shine, Koyejo, et al., 2016). Our findings, in which arousal-dependent switching of this network was not found at rest, may be consistent with the notion that central executive network switching relies on cognitive engagement, which may be deeply rooted with its role in several cognitive functions (Corbetta & Shulman, 2002).

A second goal of our study was to identify which characteristics of the fMRI time series contribute to the sensitivity of network switching to arousal state. To address this, we constructed four null models that preserved specific features of the empirical fMRI time series. For both datasets examined, we failed to reject the fourth null model, which preserved the static correlation among the salience, default mode, and central executive networks. However, the arousal-dependent network switching could not be fully explained by fMRI signal characteristics maintained in the other three null models (spatial and temporal mean and variance, and correlation with the global signal). This indicates that static correlation among these three networks is the primary characteristic driving the arousal dependency of network flexibility. Preserving this static correlation increases the likelihood of observing larger effect sizes in network switching differences across arousal states. While static correlation contributes to the arousal dependency of network switching, arousal did not moderate its relationship to participants’ task performance. Therefore, switching rate may capture neural dynamics linking arousal and cognition that go beyond these basic functional connectivity signatures.

Further, our investigation revealed that the switching rates of several parcels within the salience, central executive, and default mode networks are sensitive to arousal state. Significant parcels included the frontal regions, thalamus, insula, anterior cingulum, hippocampus and cerebellum. Notably, the thalamus parcels corresponding to all three networks exhibited a stronger arousal-dependence compared to null models that preserved the nodal mean, variance, and correlation with the global signal (null model 3). The thalamic node for the right central executive network survived all null models, revealing that it conveys unique information about brain arousal beyond basic fMRI signal characteristics. In the ascending arousal system, brainstem arousal nuclei project their signals to the thalamus before distributing to the cortex (Jones, 2003), and this pathway is essential for higher cognitive functions (Aston-Jones & Cohen, 2005). Consistent with this pathway, our findings of more frequent switching of the thalamus in the alert state may reflect its role in coordinating the activity of large-scale brain networks supporting optimal cognitive performance (Shine, Bissett, et al., 2016b).

In another avenue, we sought to determine whether the fraction of time the salience network shares its community assignment (shared allegiance) with the central executive network or the default mode network depends on arousal state. We found that the salience network showed stronger allegiance with both the central executive and default mode networks during drowsy states. One possible explanation is the well-established observation that functional connectivity across the brain increases during drowsy states (Chang et al., 2013, 2016; Liu & Falahpour, 2020), which may account for the increased prevalence of shared community assignments across these networks. Yet, this effect was not uniformly observed, as certain network pairs exhibited higher allegiance during alert states compared to drowsy states (e.g., PSAL-LCEN; **Figure 4(c,d)**). Future studies should investigate whether these patterns change during task engagement.

Lastly, we investigated whether arousal state influences how network switching relates to task performance. Prior work (Pedersen et al., 2018) showed that global brain switching predicts working memory and relational task accuracy in the HCP-3T dataset. Therefore, we conducted an exploratory analysis to test whether arousal state moderates the relationship between global brain network switching and behavioral performance on both of these cognitive tasks. We found that arousal moderated the association between global brain switching and relational task accuracy. Specifically, for participants who were drowsy during their HCP-7T scan, higher global switching was related to lower accuracy on the task. In contrast, for participants who were in an alert state, higher global switching related to better task performance. These findings provide preliminary evidence that global brain switching may serve as a neural marker of arousal-dependent cognitive changes. One limitation of this analysis, however, is that the fMRI scans and cognitive assessments were conducted months apart. We view this approach as analogous to relating resting-state fMRI measures to behavioral assessments collected outside the scanner, a prevalent practice in the study of brain-behavior relationships. A key strength of this design is that the behavioral measure was acquired in an environment similar to that of the arousal measurements. Future studies should investigate whether these results are replicated when the time between scanning and behavioral testing is shorter.

Our study has implications for psychiatric disorders in which interactions among the salience, central executive, and default mode networks are known to be abnormal and contribute to cognitive impairment (Chand et al., 2017; Menon, 2011; Suttkus et al., 2021). In particular, our findings lead us to speculate that arousal dysregulation may contribute to disruptions in the interactions among these networks, which in turn may play a role in patients’ cognitive impairments. Indeed, current investigations in this direction are emerging (Neal et al., 2023; Xie et al., 2024), but further studies in clinical populations should verify this possibility.

To conclude, we provide evidence that the dynamic activity of switching is sensitive to arousal state. While this finding suggests that arousal state may be a potential confound in previous studies identifying network switching as a marker of cognition, we interpret these results as indicating that network switching is a neural marker of arousal-dependent cognitive changes. In support of this alternative hypothesis, we provide preliminary evidence that arousal state moderates the relationship between global brain switching and performance on specific cognitive tasks.

## Online Methods

### Data 1 (HCP-7T)

#### Subjects and data acquisition

A total of 246 resting-state fMRI sessions from 133 subjects (Female=73, Male=60, ages 22-37) were included from the Human Connectome Project 7T (HCP-7T) dataset based on the following criteria: the fMRI sessions contained simultaneously recorded eye-tacking data and were classified into an alert or drowsy arousal state (see “Eye Tracking Data Pre-processing” and “Arousal Staging” sections). All subjects provided written informed consent, and human subject protocols were approved by the University of Minnesota Institutional Review Board. Resting-state fMRI data was acquired (flip angle = 45 deg, TR = 1000 ms, echo time = 22.2 ms, voxel size = 1.6 × 1.6 × 1.6 mm3, matrix size = 130×130, 85 axial slices, acceleration factor = 2) in four runs of 15 minutes each (900 volumes). At the beginning of each of the four 7T imaging sessions, participants had their eyes open with relaxed fixation on a bright cross-hair projected on a dark background. Among the four sessions, oblique axial acquisitions alternated between phase encoding in a posterior-to-anterior (PA) direction in runs 1 and 3, and an anterior-to-posterior (AP) phase encoding direction in runs 2 and 4. These subjects also underwent task fMRI sessions in a 3T scanner (HCP-3T dataset), which included working memory and relational tasks.

#### fMRI preprocessing

Full details of the data acquisition and preprocessing are provided elsewhere (Glasser et al., 2013; Jenkinson et al., 2002, 2012). In brief, fMRI data was collected using a Siemens 7T Magnetom actively shielded scanner and a 32-channel receiver coil array with a single channel transmit coil (Nova Medical, Wilmington, MA). Whole-brain fMRI data was processed using the HCP pipelines (Glasser et al., 2013), which corrects for head motion and EPI spatial distortion and brings the fMRI data into alignment with the MNI standard space, as described in Benson et al., 2018. The fMRI data were also denoised for spatially specific structured noise using multirun spatial ICA and FIX (Glasser et al., 2018).

#### Eye Tracking Data Pre-processing

We followed the procedures from Gonzalez-Castillo et al., 2022 to preprocess the eye-tracking data. Briefly, we removed pupil-size data that occurred before the onset of each fMRI scan in order to synchronize the eye-tracking data to the fMRI data. We then linearly interpolated the pupil size over blink events that lasted for less than one second. Blink events that lasted longer than one second were considered eye-closure periods and were not interpolated. To improve identification of eye closures, we zeroed out short-duration (<100 ms) bursts of eyes-open periods that occurred in between long-duration blink events. (Gonzalez-Castillo et al., 2022).

#### Arousal Staging

The preprocessed eye-tracking data were used to classify each scan into an alert or drowsy arousal state based on the procedure of Gonzalez-Castillo et al., 2022.

*a*.) *Alert Scans* were defined as scans that had eye closures for less than 5% of the scan duration.
*b*.) *Drowsy scans* were defined as scans that had eye closures for a range of 50% to 90% of the scan duration.

Scans with eye closures greater than 90% of the time were discarded to exclude potentially faulty eye-tracking data and to exclude sessions where subjects may not have complied with the eyes-open instructions. Scans with eye closures between 5-50% were considered as an intermediate arousal stage (not clearly drowsy or alert) and excluded from the analysis. We implemented a more stringent threshold for the drowsy state classification compared to Gonzalez-Castillo et al., 2022 (50% instead of 20%) because we aimed to include sessions that could be more confidently assigned as alert or drowsy.

#### Working Memory Task and Relational Task

Task performance measures used in this analysis were drawn from the HCP-3T task fMRI dataset. Details of these tasks are described in Barch et al., 2013. In summary, the working memory task was a N-back task that presented 4 types of stimuli that were either pictures of faces, places, tools, or body parts that differed across each block. In each run of the task, there were a total of 8 blocks: 4 blocks corresponded to the 0-back condition, where participants were instructed to simply press a button when a stimulus appeared, and the other 4 blocks corresponded to the 2-back condition, where participants were instructed to determine if the stimulus presented in the current trial was also presented two trials previously. The relational task was also administered in the scanner and measured participants’ ability to form relational matches, defined as the ability to identify similarities or differences between pairs of objects. Relational matching was assessed by asking participants to determine what was similar or different (shape or texture) among four objects presented on the screen (two on the top and two on the bottom) (Barch et al., 2013; Smith et al., 2007).

### Dataset 2 (VU-EEG-fMRI)

#### Subjects and data acquisition

We included 18 subjects (22 scans, nine females, aged 31 ± 13 years) in this study, each of whom underwent at least one simultaneous EEG-fMRI scan that passed quality control for motion artifacts and maintained either an alert or drowsy state across the entire scan. All subjects provided written informed consent and human subjects protocols were approved by the Institutional Review Board of Vanderbilt University.

MRI data was acquired on a 3T Elition X scanner (Philips Healthcare, Best, Netherlands) with a 32-channel head/neck coil. A high-resolution, T1-weighted structural image (TR = 9 ms, TE = 4.6 ms, flip angle = 8 deg, matrix = 256×256, 150 sagittal slices, 1 mm isotropic) was acquired for anatomic reference. A multi-echo EPI sequence was used to acquire the resting-state fMRI scans (flip angle = 79 deg, TR = 2100 ms, TE = 13.0, 31.0, and 49.0 ms, voxel size = 3 × 3 × 3 mm3, slice gap = 1 mm), and subjects were instructed to keep their eyes closed. One or two scans of 20-minutes duration (575 volumes) were acquired for each subject. Scalp EEG was acquired simultaneously with fMRI using a 32-channel MR-compatible system (BrainAmps MR, Brain Products GmbH) at a sampling rate of 5 kHz, and was synchronized to the MRI scanner’s 10 MHz clock to facilitate reduction of MR gradient artifacts. The EEG data were referenced to the FCz channel. Photoplethysmography (PPG) and respiration belt signals were also acquired during the scans (Philips). The PPG transducer was placed on the left index finger, and MRI scanner (volume) triggers were recorded together with the physiological and EEG signals for data synchronization.

#### fMRI preprocessing

The resting-state fMRI scans were processed following the methods described in Pourmotabbed et al., 2025. In summary, motion co-registration and slice-timing correction were carried out using the functions *3dvolreg* and *3dTshift* in AFNI. For motion co-registration, the alignment parameters were estimated only for the time series of the second (middle) echo, and the resulting parameters were applied to the time series of all three echoes. Following this initial processing, multi-echo ICA denoising was carried out using *tedana* 0.0.9a (DuPre et al., 2020; Kundu et al., 2012, 2013). The fMRI data were then nonlinearly registered to a standard-space MNI152 template using ANTS, followed by spatial smoothing (FWHM 3 mm) and fourth-order polynomial detrending in AFNI.

#### EEG preprocessing

The EEG data was processed in the following manner: reduction of gradient and ballistocardiogram (BCG) artifacts was carried out using BrainVision Analyzer 2 (Brain Products, Munich, Germany) (Moehlman et al., 2019). Gradient artifact reduction followed the average artifact subtraction technique (Allen et al., 2000). BCG artifact correction was carried out by subtracting an artifact template locked to cardiac R-peaks detected from the EKG channel, after accounting for an estimated temporal offset between the R-peak and the BCG artifact. Following gradient and BCG artifact correction, EEG data were temporally aligned to the fMRI data and down sampled to 250 Hz.

#### Arousal Staging

Arousal states were classified using an EEG automated arousal staging program, Vigilance Algorithm Leipzig (VIGALL) (Ulke et al., 2017) and were validated by visual inspection of the EEG spectrograms (De Gennaro et al., 2001; Marzano et al., 2011; Tsuno et al., 2002). VIGALL has been used extensively in an array of studies to characterize arousal level in various psychiatric disorders (Kramer et al., 2019; Olbrich et al., 2013; Ulke et al., 2019). Its ability to track arousal states has also been validated against heart rate and skin conductance recordings (Huang et al., 2015). The VIGALL algorithm was used to stage each 1-second time window of the EEG data into five discrete arousal states (A1, A2, A3, B1, B2/3) after interpolating missing channels in the VIGALL standard and re-referencing to the common average (see Pourmotabbed et al., 2025). This approach yielded a temporal arousal rank that fluctuated between 2 and 6 across time. A scan was classified as *drowsy* if the average arousal level across the scan was below 3.5 and as *alert* if it exceeded 3.5. We also identified scans in which arousal ranks were initially high (4–6) but declined to lower levels (2–3) by the end of the scan; these were labeled as *transition* scans. However, the number of transition scans was insufficient to support within-subject analyses. Therefore, for the present study, we included only scans that were consistently classified as either alert or drowsy for the entire duration of the scan. We validated our findings by inspecting each scan’s EEG spectrogram, ensuring that alert scans had prominent alpha-band power throughout the entire scan; for drowsy scans, we ensured they had a prominent theta– and/or delta-band power throughout the scan.

### Deriving Measures and Analysis

#### Deriving Large Scale Networks

Large scale networks were derived using a priori atlas (“FINDLAB atlas”) which contains 14 pre-defined networks derived on a separate fMRI dataset (Shirer et al., 2012). This atlas contains the following networks: anterior salience network (ASAL), posterior salience network (PSAL), auditory network (AUD), basal ganglia (BasG), dorsal default mode network (DDMN), ventral default mode network (VDMN), high-level visual network (HVIS), language network (LANG), left central executive network (LCEN), right central executive network (RCEN), precuneus (PREC), primary visual network (PVIS), sensorimotor (SMOTOR), and visuospatial network (VISSPAT). To derive each network’s time series, we performed dual regression (Nickerson et al., 2017). Dual regression is a two-step procedure in which spatial brain templates (here, atlas-defined network masks) are first spatially regressed onto individual fMRI scans on a volume-by-volume basis to estimate subject-specific time series, and these time series are then temporally regressed back onto the data (specifically, onto the time series of each voxel) to obtain subject-specific spatial maps. For the parcel analysis, we used the FINDLAB atlas 90-ROI parcellation of the 14 pre-defined networks and took the mean of the voxel-wise signals within each ROI to derive the parcel time series. For defining the regional labels of the parcels, we mapped regions to the Automated Anatomical Labeling (AAL) 3 atlas (Rolls et al., 2020) via visual inspection.

#### Computing Network Switching and Global Network Switching

Network switching is quantified using multilayer modularity. Traditionally, modularity quantifies the extent to which nodes in a network segregate into densely interconnected communities. Multilayer modularity extends this approach by jointly estimating community structure across successive time windows while preserving temporal consistency of module identity, thereby enabling the quantification of network switching. To compute network switching, we adapted the methods described in Pedersen et al., 2018. The first step involves computing the sliding-window Pearson correlation between each pair of nodes and setting any negative correlations to 0. For our investigation, the window lengths for both VU-EEG-fMRI and HCP-7T datasets were approximately three minutes, and the overlap between successive sliding windows was 1 TR. Next, we used an iterative ordinal Louvain algorithm to optimize network modularity across windows (Mucha et al., 2010). [implemented with software by Lucas G. S. Jeub, Marya Bazzi, Inderjit S. Jutla, and Peter J. Mucha, “A generalized Louvain method for community detection implemented in MATLAB,” netwiki.amath.unc.edu/GenLouvain/GenLouvain]. Multilayer modularity was estimated by maximizing the modularity quality function Q, which identifies community assignments across sliding windows while accounting for temporal coupling between layers. The modularity Q was computed using the following equation:

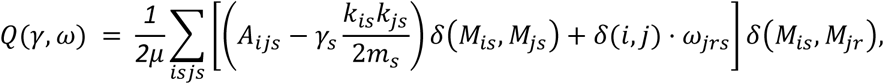

Here, A_ijs_ is the sliding-window dynamic correlation matrix between node (either parcel or network) (i) and (j) for time point (s). k_is_ k_js_/ 2ms is a ratio between node connectivity value (k) for node (i) and (j) at TR (s), and m is the sum of all node connectivity values. γ_s_ is the topological resolution parameter of time point and controls for the size of community, and ω_jrs_ is the temporal coupling parameter that measures the strength for node j between time window r and s. δ(*M*_*is*_, *M*_*js*_) *and* δ(*M*_*is*_, *M*_*jr*_) are used to control temporal coupling: when nodes j and i are in the same community/ module, this value is set to 1; and if they are in a different module, it is set to 0. To improve stability in this investigation, we ran multilayer modularity on a consensus matrix rather than on the raw fMRI dynamic connectivity windows. The consensus matrix was constructed by first running multilayer modularity 10 times on fMRI dynamic connectivity windows and storing the community assignments of each iteration, and then computing the percentage of times for which each pair of nodes was assigned to the same community. Subsequently, we used the code “flexibility” (that can be retrieved from http://commdetect.weebly.com/) to calculate the percentage of time-windows across a scan for which each network switches into a new community assignment, which is what we term *switching* (Bassett et al., 2011b).

We computed switching at both the network and parcel levels. For the network-level analysis, nodes corresponded to the 14 predefined subnetworks from the FINDLAB atlas, so switching reflected the amount of time an individual subnetwork switched between communities formed between the 14 subnetworks. A measure of global brain switching was defined by averaging the switching rates of the individual subnetworks. For the parcel-level analysis, nodes consisted of the 90 ROIs (parcels) defined by the FINDLAB atlas. Parcel-level switching reflects the amount of time each parcel switched across the communities formed from these 90 ROIs.

### Null models

To identify which characteristics of the BOLD signal may contribute to arousal-dependent network flexibility characteristics, we constructed four null models. For each model, we generated 100 new (surrogate) time series per scan that maintained certain parameters from the empirical data, specifically: (model 1) mean across time and across nodes (either at network or parcel level); (model 2) mean across time and across nodes, and variance across time and across nodes; (model 3) mean across time and across nodes, variance across time and across nodes, and the correlation between each node and the global signal; and lastly, (model 4) mean across time and across nodes, variation across time and across nodes, the correlation between each node and the global signal, and the static correlation between the salience, default and central executive subnetworks. After obtaining the 100 surrogate time series from a given null model, we calculated how many times the effect size (here, for the difference in switching rates between arousal states) of this “null” data exceeded the effect size of the empirical data, yielding a p-value. All four models were tested for the VU-EEG-fMRI dataset. The null model tests were conducted for the EEG-fMRI dataset first, and after determining that most networks of interest survived null model 1 and 2, we proceeded to test only null models 3 and 4 for the HCP-7T data for computational efficiency. Given the larger size of the HCP-7T data, here we constructed 10 surrogate time series for each of the 246 scans.

#### Community assignment comparison across arousal state

The multilayer modularity analysis yields a matrix of each node’s community assignment across each sliding window. From this matrix, we quantified the fraction of time windows in which two networks were in the same community. We computed this fraction for each scan, and defined this as the strength of community allegiance between a given pair of networks. Next, we tested for differences in community allegiance across arousal states (alert vs drowsy). We hypothesized that the amount of time in which the salience network would be in the same community with the central executive network would be higher in alert states compared to drowsy states. In contrast, we hypothesized that the amount of time in which the salience network would be in the same community with the default mode network would be higher in drowsy states compared to alert states. To test these hypotheses, we conducted Mann-Whitney U tests for the VU-EEG-fMRI data and the HCP-7T data. The percentage of shared community assignments for all networks is provided in **Supplementary Figure 3**.

#### Comparing static network and global correlations across arousal state

We used a Mann-Whitney U test to examine whether the correlation between ASAL, DDMN, and VDMN with the global signal differed across arousal states. Additionally, we tested if the static correlation, computed by taking the average correlation across all 14 sub networks, was altered between arousal states (again using a Mann-Whitney U test).

#### Global brain switching across arousal state and as a moderator of task accuracy

To test if global network switching is altered across arousal states, we conducted Mann-Whitney U tests for both the VU-EEG-fMRI data and the HCP-7T data (**Figure 1**). To test if arousal state moderates how global brain switching relates to two selected task accuracy measures, we computed global network flexibility as described above, followed by a moderation test. Specifically, we constructed a linear model where we tested if global switching and arousal state predicted working memory or relational task accuracy while testing for an interaction/moderation (collected in the HCP-3T data).

## Data and Code Availability Statement

This study used data acquired from Vanderbilt University and is in process of being publicly available. If interested in acquiring this data, please reach out to Dr. Catie Chang at. The Human Connectome 7T data is publicly available at https://www.humanconnectome.org/study/hcp-young-adult/data-releases. Code for reproducing this study will be available on github.com/neurdylab/netswitch_across_arousal and preliminary code is currently available on https://github.com/krogge-obando/net_switching_across_arousal.

## Acknowledgments

Data were provided in part by the Human Connectome Project, WU-Minn Consortium (Principal Investigators: David Van Essen and Kamil Ugurbil; 1U54MH091657) funded by the 16 NIH Institutes and Centers that support the NIH Blueprint for Neuroscience Research; and by the McDonnell Center for Systems Neuroscience at Washington University. The authors would also like to acknowledge Dr. Allison Leich Hilbun for consultation on statistical analysis and Anastaczja Cumberjack for consultation on writing. Parts of this work were presented in abstract form at the 2023 meetings of the Organization for Human Brain Mapping, the Society for Biological Psychiatry, and Society for Neuroscience.

## Author contributions

KKO – conceptualization, data curation, formal analysis, investigation, methodology, project administration, visualization, Writing—original draft, Writing—review & editing; HP – data curation, – methodology, investigation; KK – data curation, – visualization, SW – conceptualization; JGL-validation; SG – data curation, methodology; CM; data curation, methodology; VM; Resources; DE-Resources; LQU – Writing-review-editing; MR – conceptualization, formal analysis, methodology, software, Writing-review-editing; CC-resources; conceptualization, writing-original draft, writing-review-editing, project administration.

## Supplementary Material

**Supplementary Table 1.**
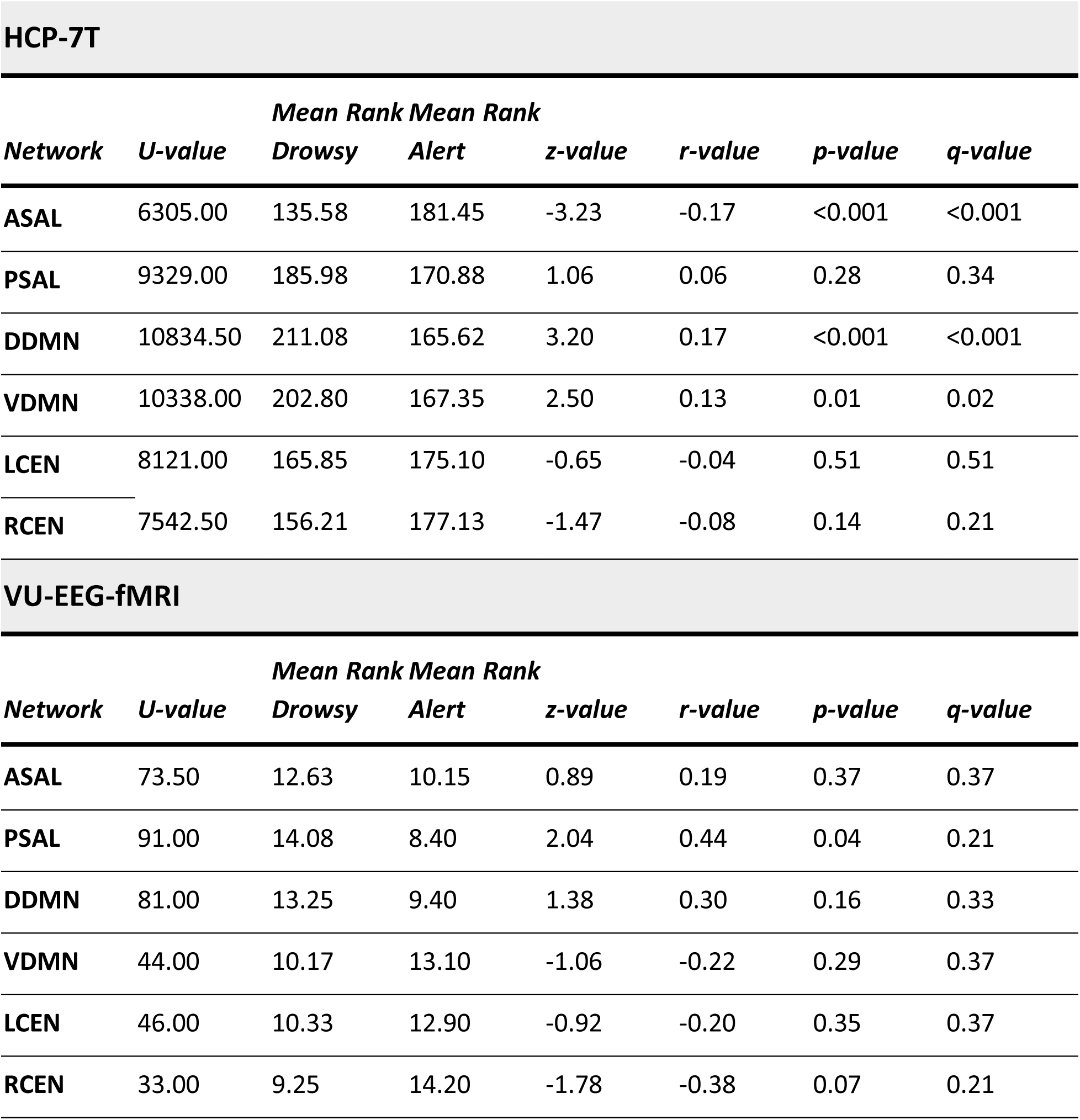
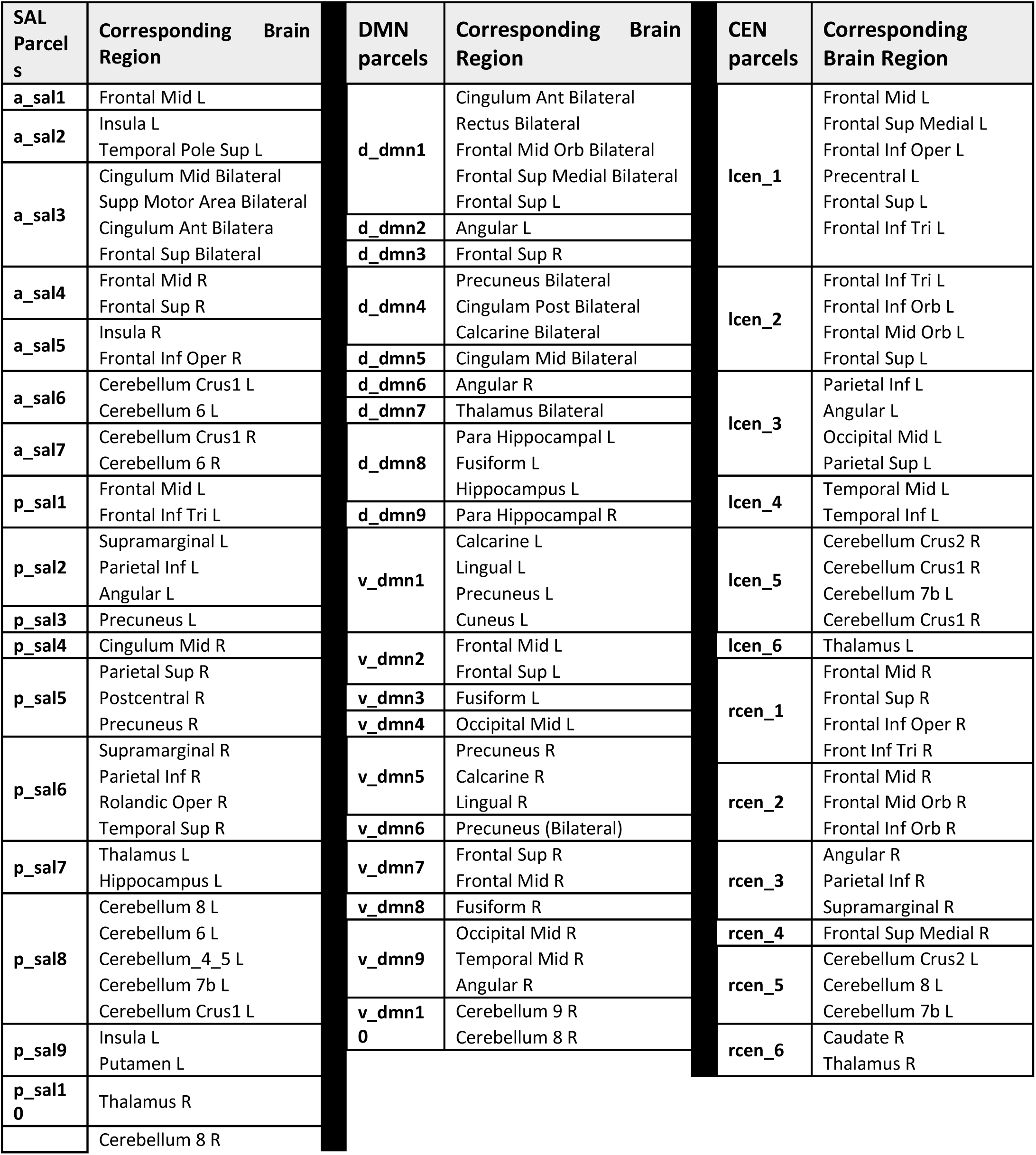
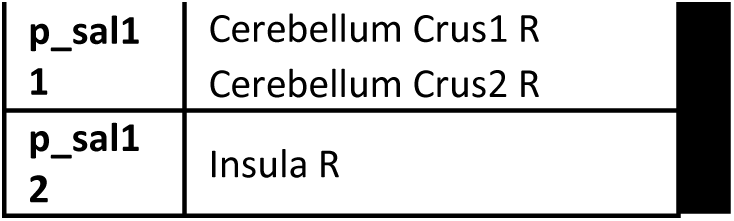
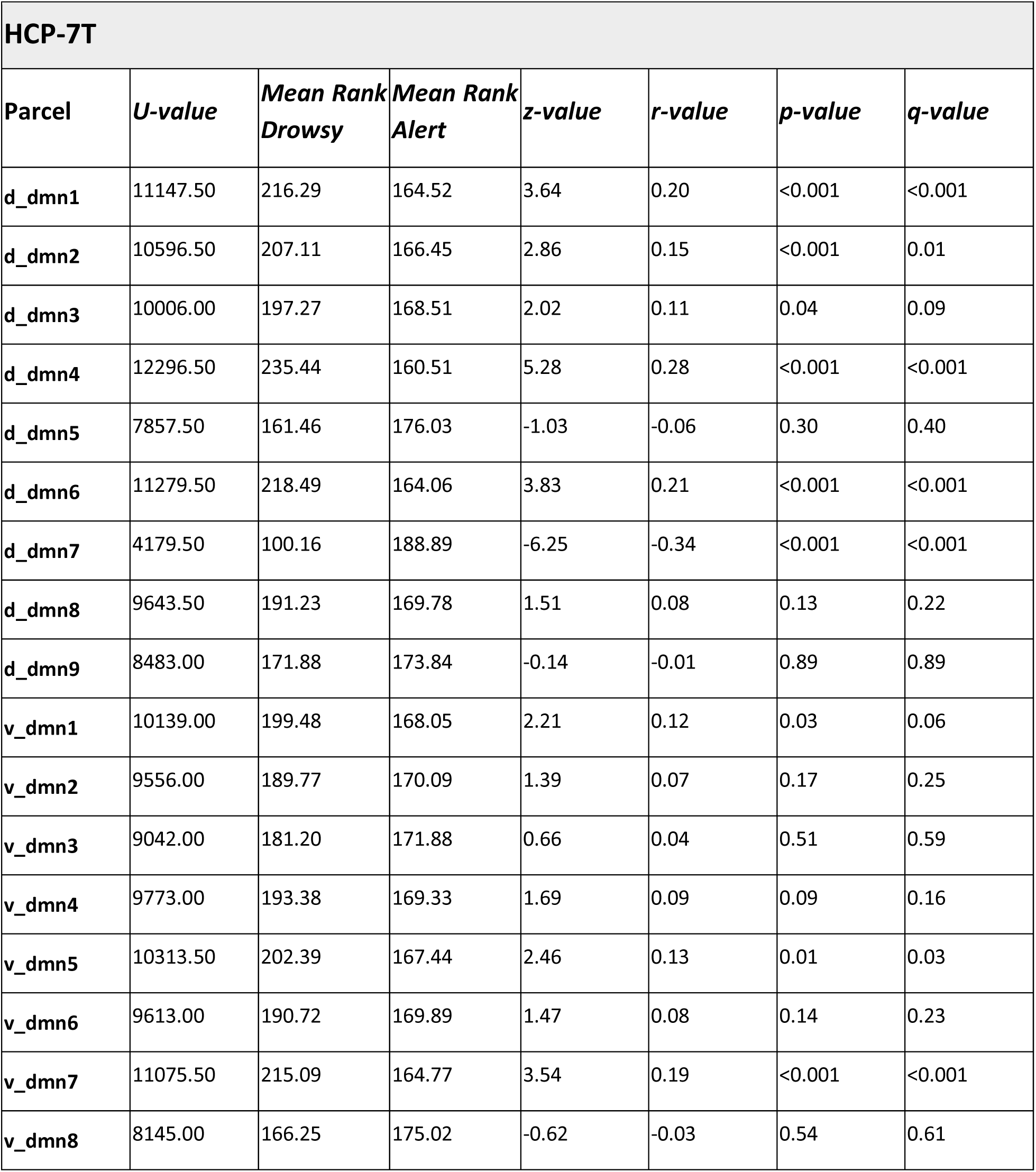
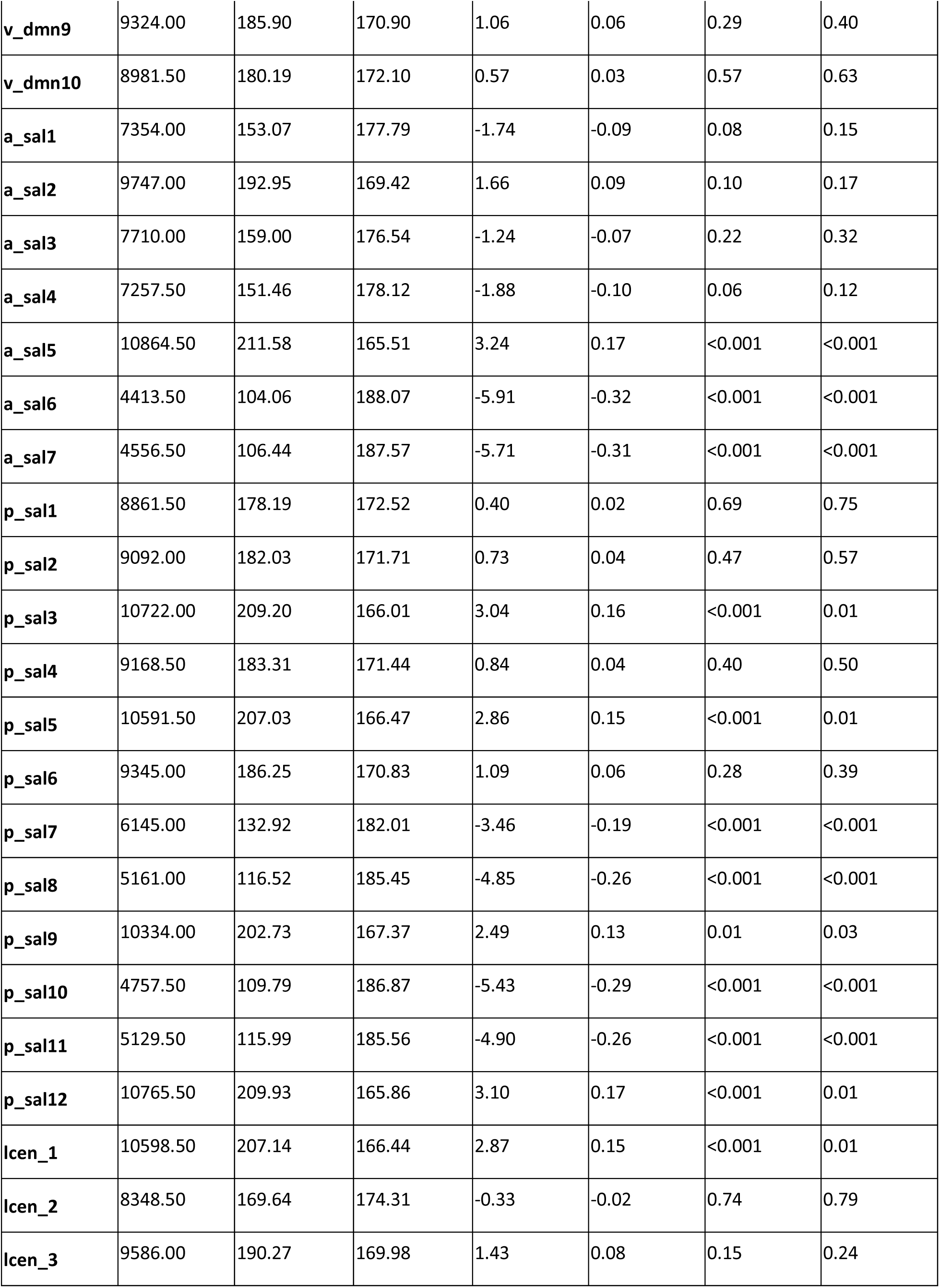
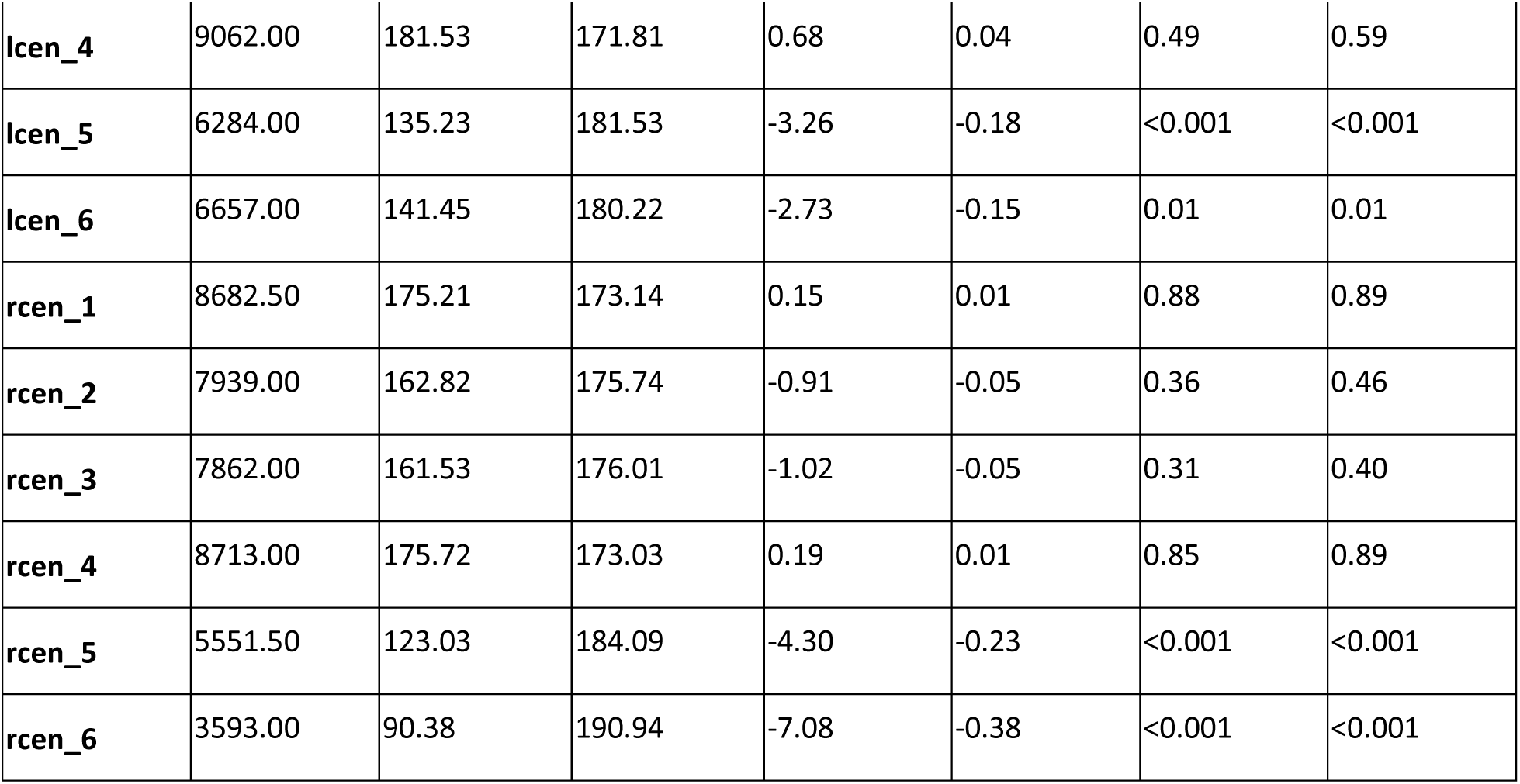
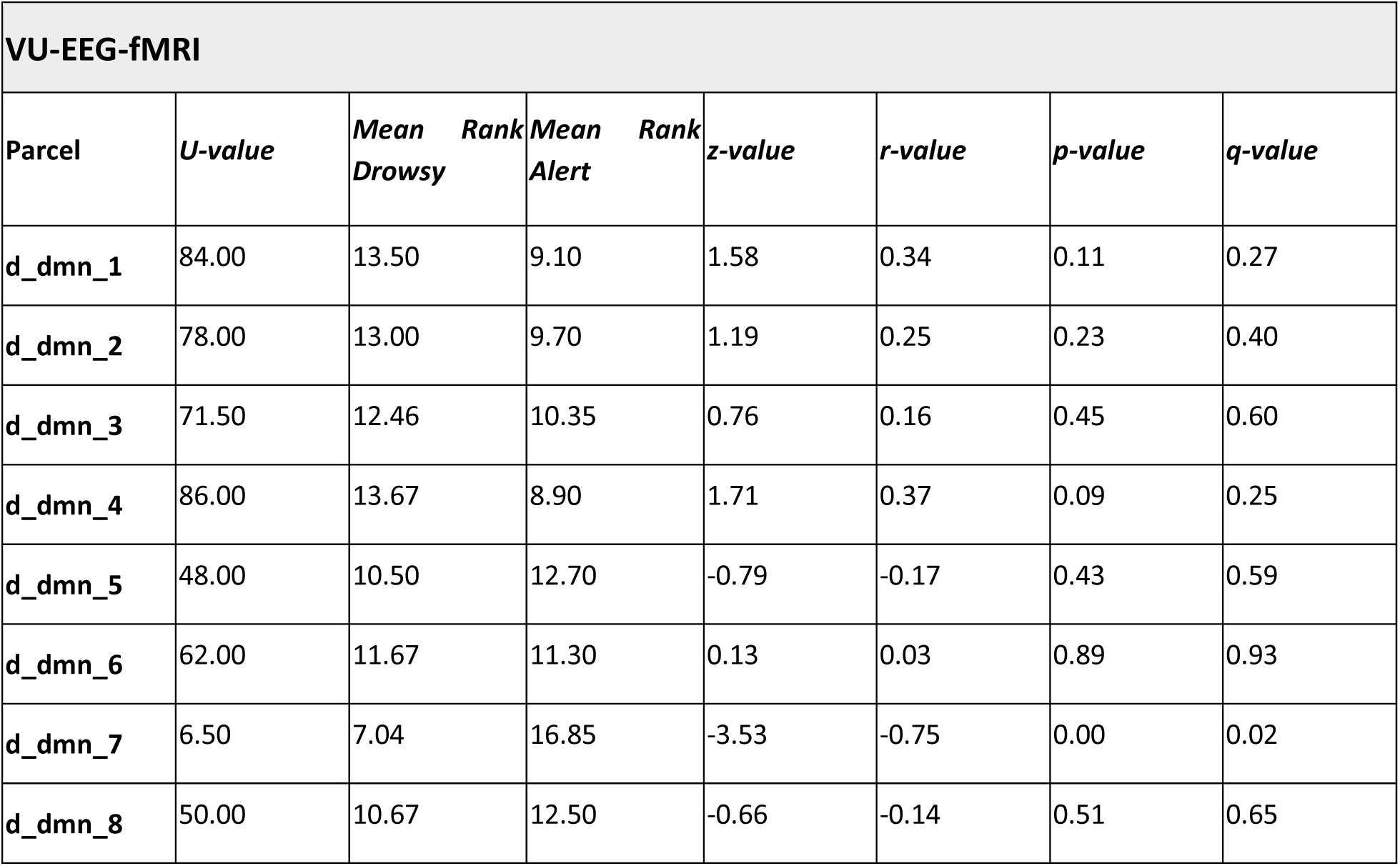
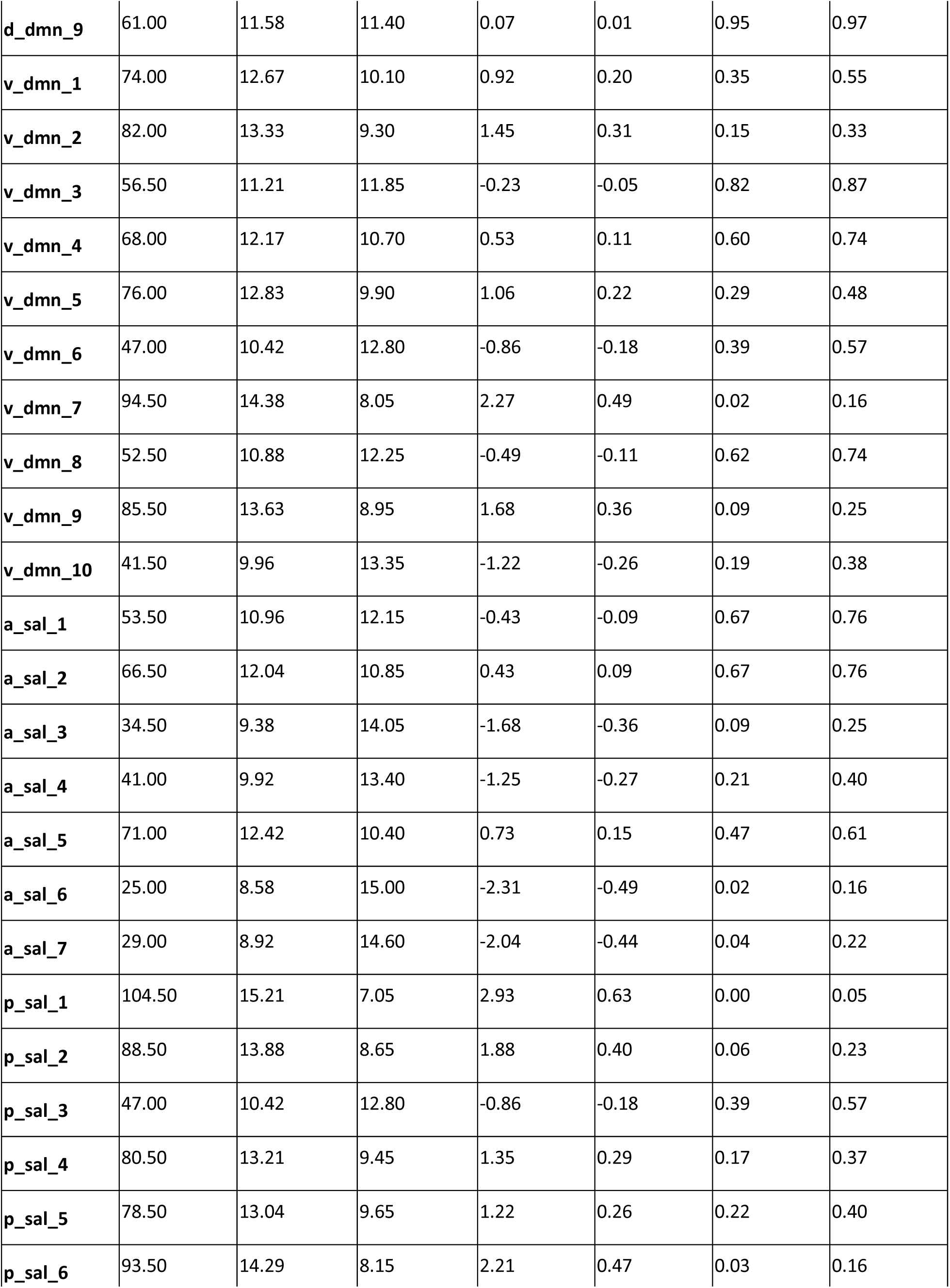
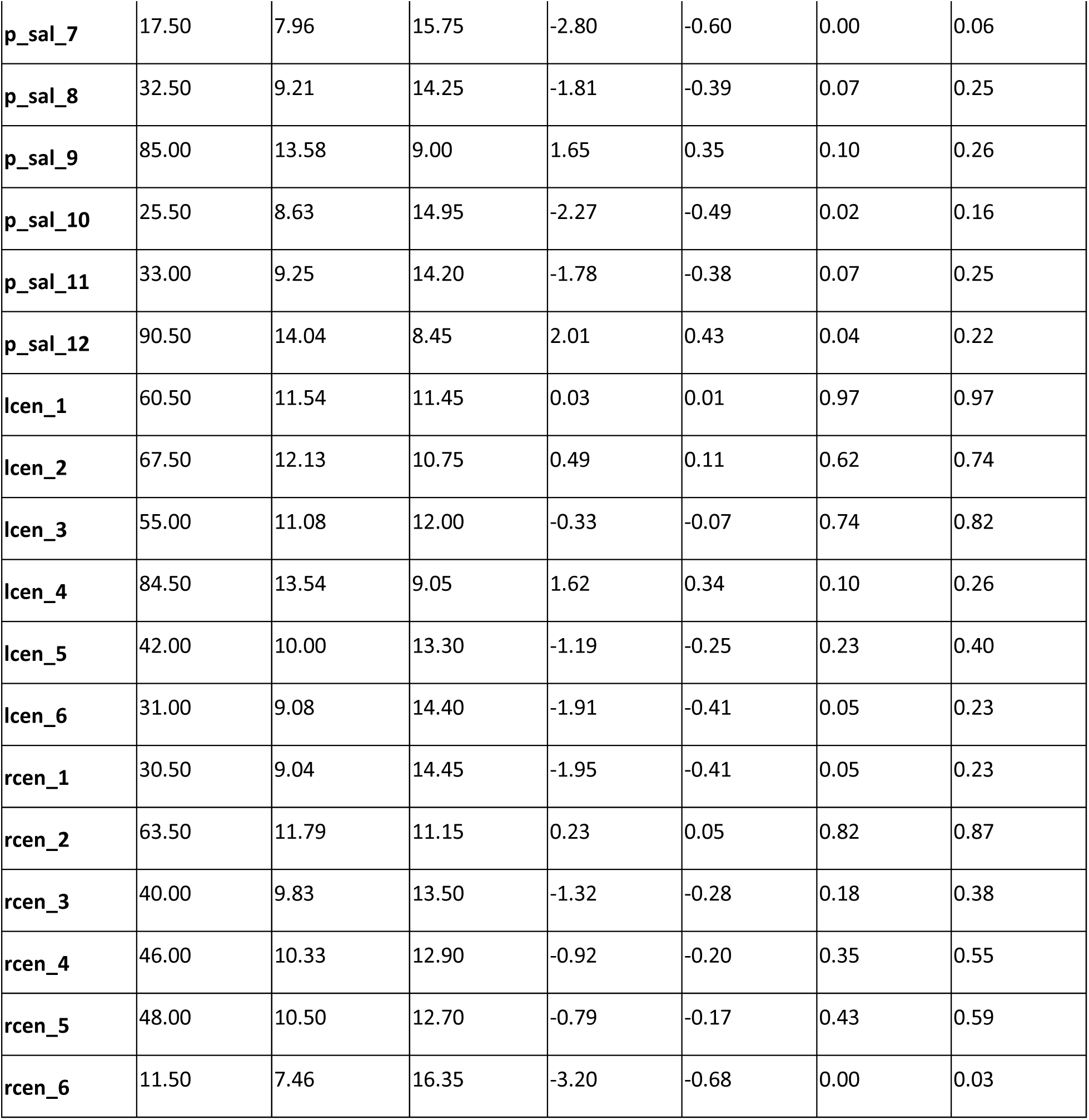
Statistical testing of arousal-dependent changes in network flexibility. Mann-Whitney U test statistics for network flexibility analyses with (a) sub-networks as nodes, or (b) label of parcels as nodes (c) statistical results for HCP-7T parcel level, (d) statistical results for VU-EEG-fMRI parcel level.

**Supplementary Table 2.**
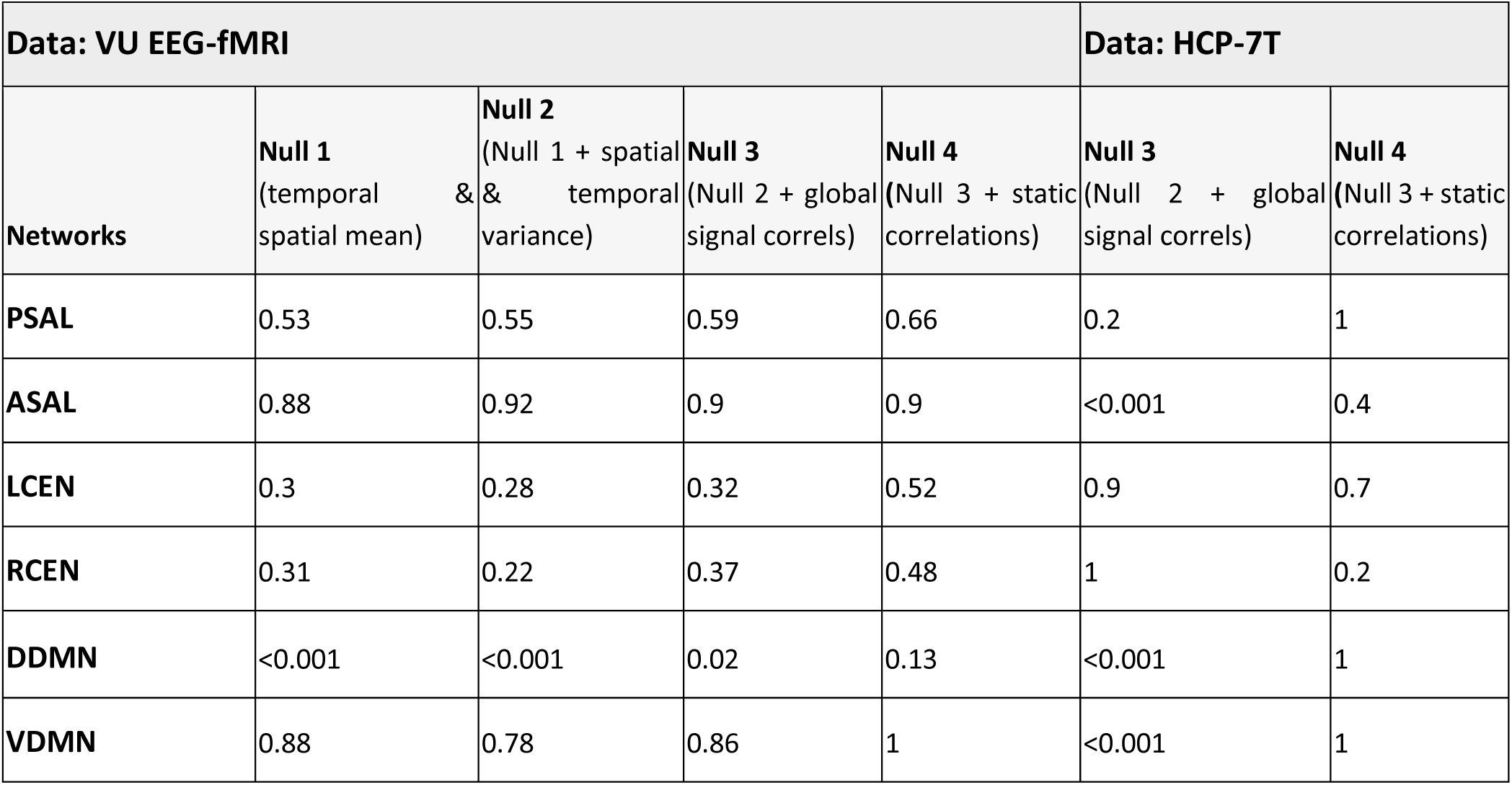
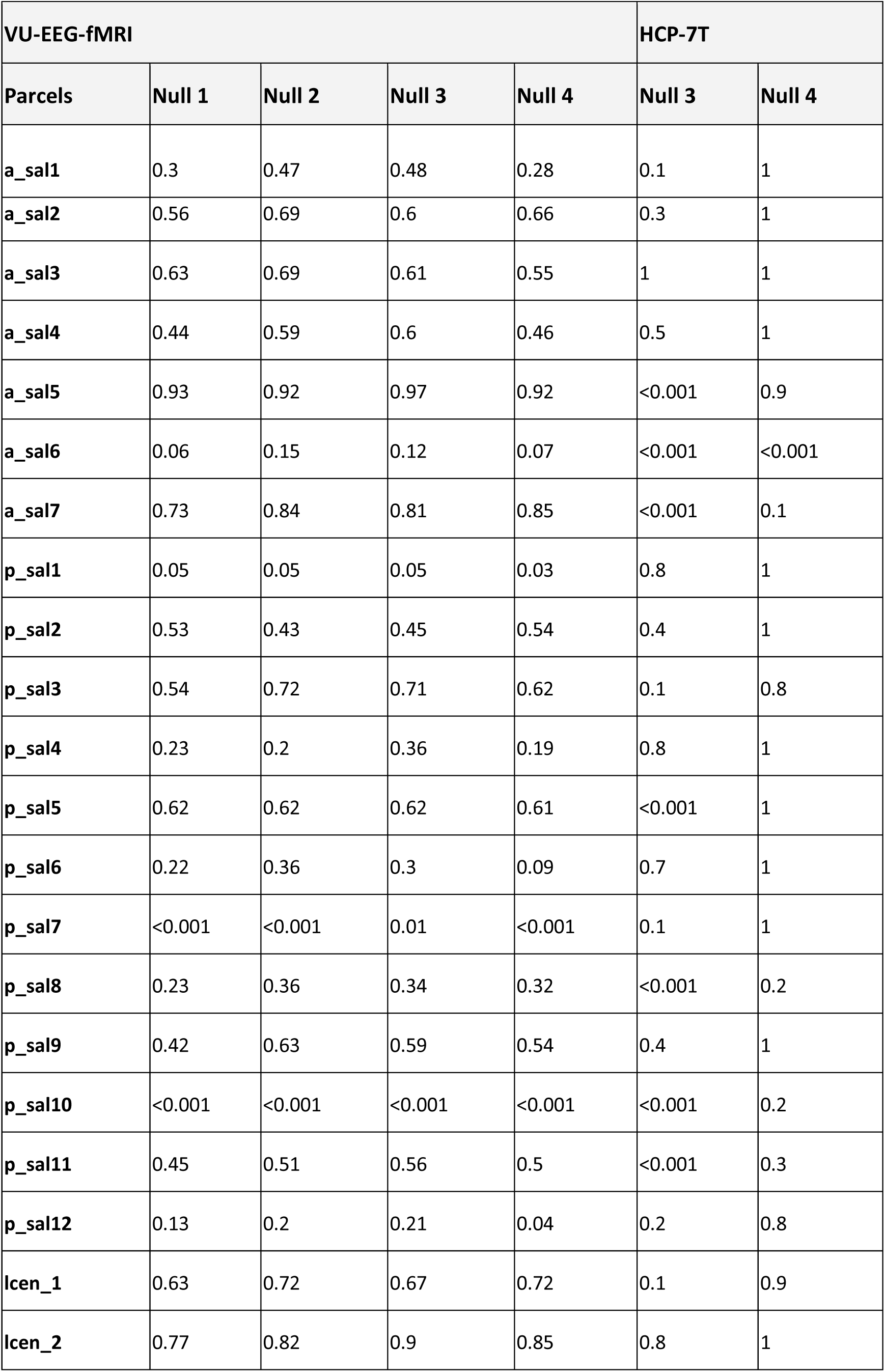
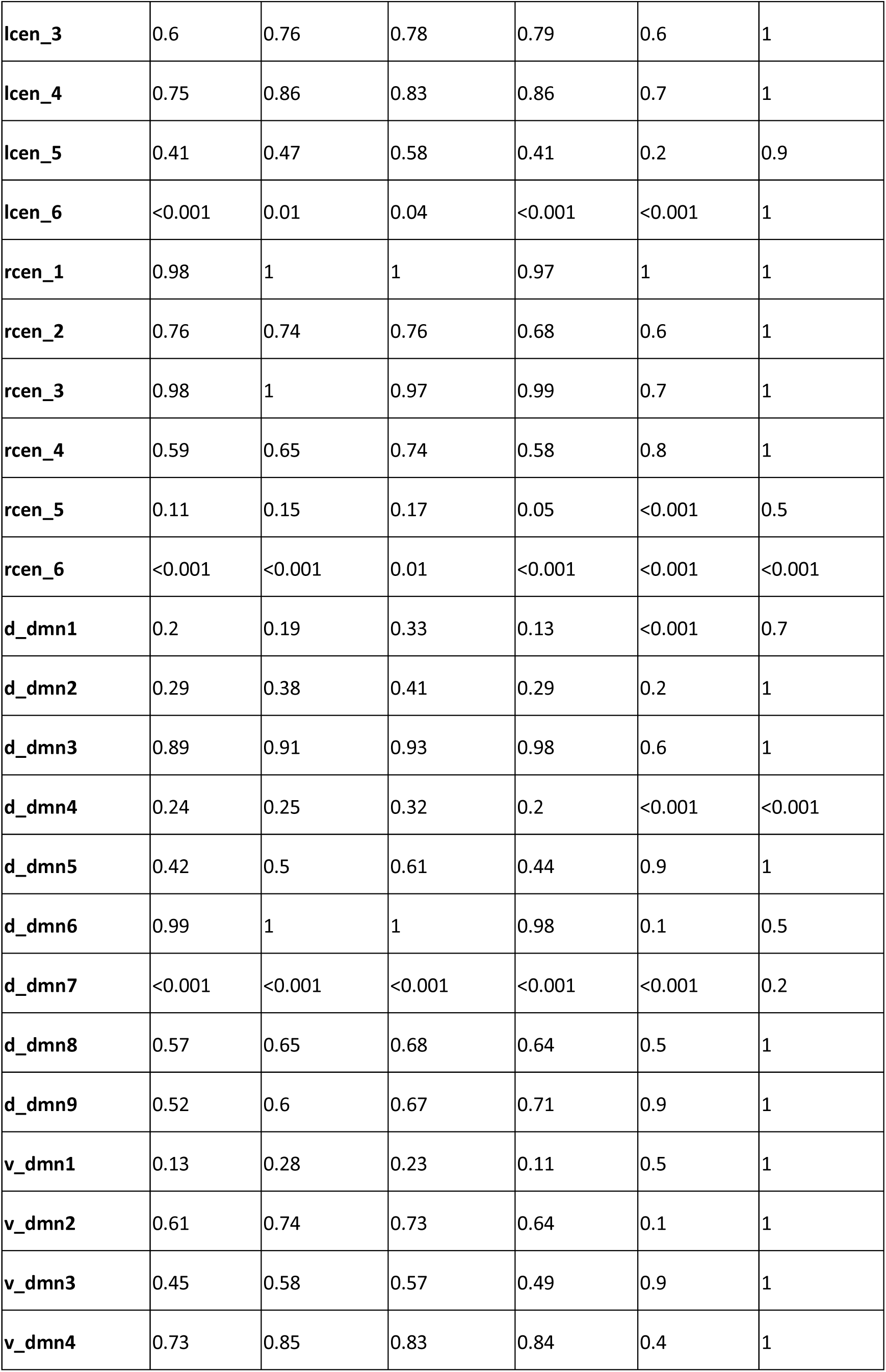
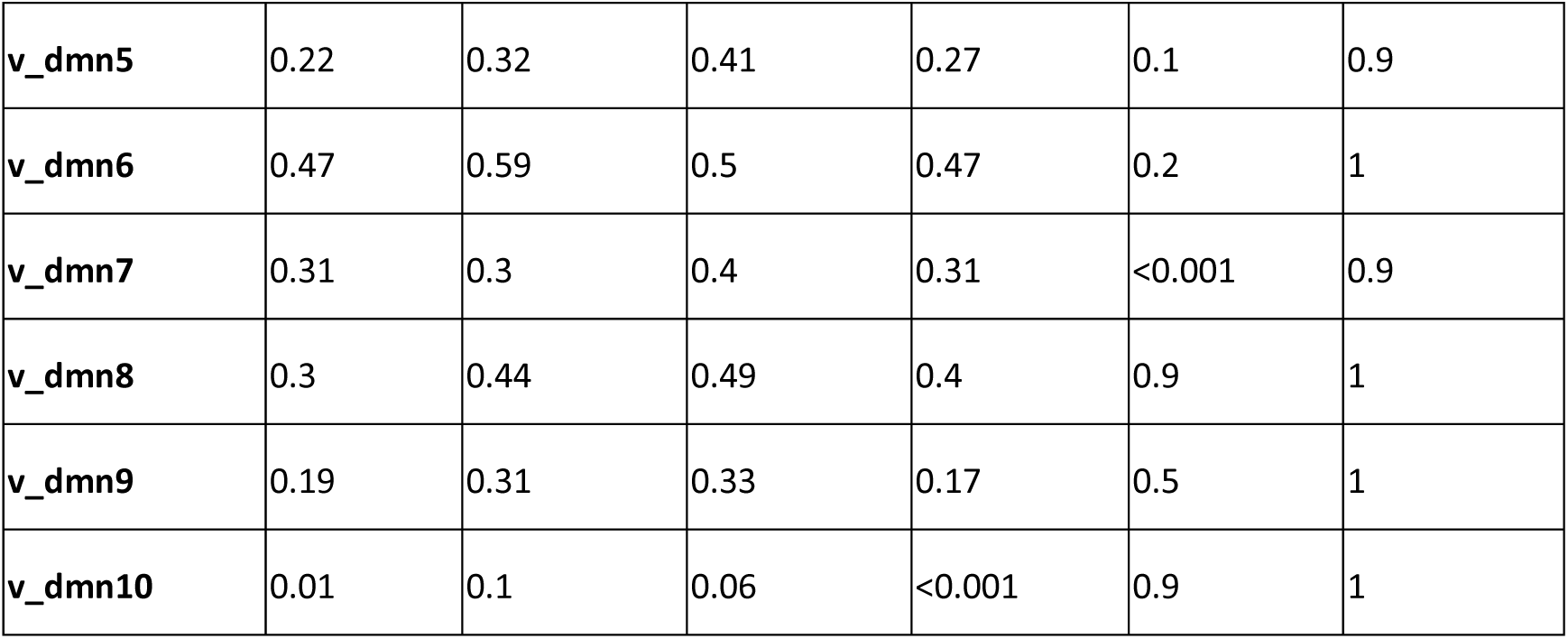
Testing which BOLD signal characteristics contribute to arousal-dependent network flexibility. a) Null model results for network-level analysis. b) Null model results for parcel-level analysis. For parcel level labels of brain regions please refer to Supplementary Table1b. Null models progressively maintained the following characteristics of the fMRI data: (1) mean across time and across nodes, (2) variation across time and across nodes, (3) the correlation between each node and the global signal, and lastly, (4) static correlation between salience, default and central executive subnetworks.

**Supplementary Figure 1.**
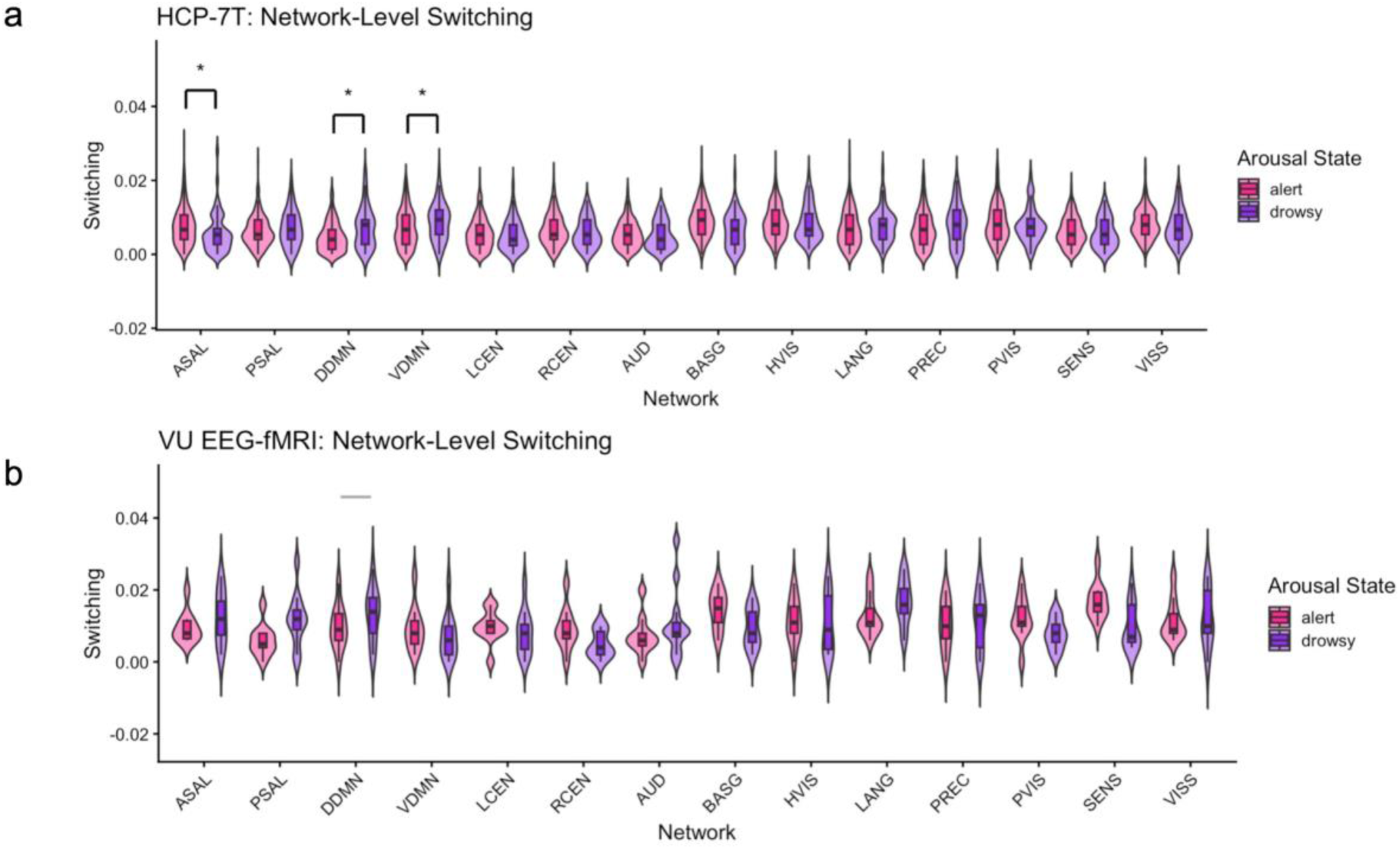
Extended network switching across arousal state. a) HCP-7T data: Violin plots show network-level switching across all 14 large-scale networks in the FINDLAB atlas for alert (pink) and drowsy (purple) arousal states. Auditory Network (AUD), Basil Ganglia Network (BASG), Higher Visual Network (HVIS), language network (LANG), precuneus network (PREC), Visual Network I (PVIS), Sensorimotor Network (SENS), Visual Network II (VISS) Note: AUD, BASG, HVIS, LANG, PREC, PVIS, SENS, VISS networks shown here were not tested for significant differences. b) Similar plot but for VU-EEG-fMRI dataset, the grey bar signifies this network was significant up to the third null model.

**Supplementary Figure 2.**
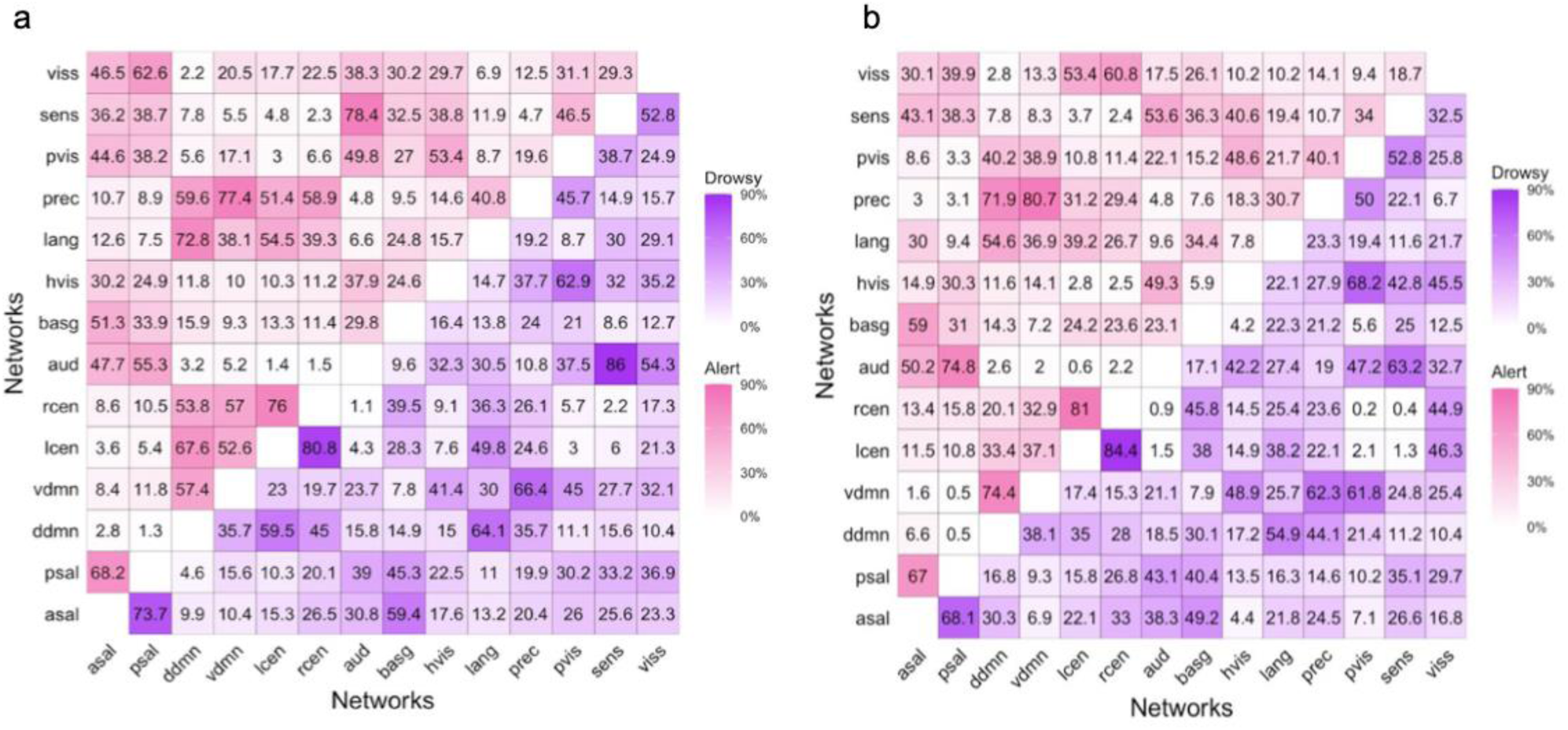
State-dependent community allegiance of all subnetworks in the FINDLAB atlas, shown for a) the HCP-7T dataset and b) the VU-EEG-fMRI dataset.

## References

1. Alameda, C., Avancini, C., Sanabria, D., Bekinschtein, T. A., Canales-Johnson, A., & Ciria, L. F. (2024). Staying in control: Characterizing the mechanisms underlying cognitive control in high and low arousal states. British Journal of Psychology, 115(4), 665–682. 10.1111/bjop.12715

2. Allen, P. J., Josephs, O., & Turner, R. (2000). A Method for Removing Imaging Artifact from Continuous EEG Recorded during Functional MRI. NeuroImage, 12(2), 230–239. 10.1006/nimg.2000.0599

3. Aston-Jones, G., & Cohen, J. D. (2005). An integrative theory of locus coeruleus-norepinephrine function: Adaptive gain and optimal performance. Annual Review of Neuroscience, 28, 403–450. 10.1146/annurev.neuro.28.061604.135709

4. Barber, A. D., John, M., DeRosse, P., Birnbaum, M. L., Lencz, T., & Malhotra, A. K. (2020). Parasympathetic arousal-related cortical activity is associated with attention during cognitive task performance. NeuroImage, 208, 116469. 10.1016/j.neuroimage.2019.116469

5. Barch, D. M., Burgess, G. C., Harms, M. P., Petersen, S. E., Schlaggar, B. L., Corbetta, M., Glasser, M. F., Curtiss, S., Dixit, S., Feldt, C., Nolan, D., Bryant, E., Hartley, T., Footer, O., Bjork, J. M., Poldrack, R., Smith, S., Johansen-Berg, H., Snyder, A. Z., & Van Essen, D. C. (2013). Function in the Human Connectome: Task-fMRI and Individual Differences in Behavior. NeuroImage, 80, 169–189. 10.1016/j.neuroimage.2013.05.033

6. Bassett, D. S., Wymbs, N. F., Porter, M. A., Mucha, P. J., Carlson, J. M., & Grafton, S. T. (2011a). Dynamic reconfiguration of human brain networks during learning. Proceedings of the National Academy of Sciences of the United States of America, 108(18), 7641–7646. 10.1073/pnas.1018985108

7. Betzel, R. F., Satterthwaite, T. D., Gold, J. I., & Bassett, D. S. (2017). Positive affect, surprise, and fatigue are correlates of network flexibility. Scientific Reports, 7(1), 520. 10.1038/s41598-017-00425-z

8. Braun, U., Schäfer, A., Walter, H., Erk, S., Romanczuk-Seiferth, N., Haddad, L., Schweiger, J. I., Grimm, O., Heinz, A., Tost, H., Meyer-Lindenberg, A., & Bassett, D. S. (2015a). Dynamic reconfiguration of frontal brain networks during executive cognition in humans. Proceedings of the National Academy of Sciences of the United States of America, 112(37), 11678–11683. 10.1073/pnas.1422487112

9. Chand, G. B., Wu, J., Hajjar, I., & Qiu, D. (2017). Interactions of the Salience Network and Its Subsystems with the Default-Mode and the Central-Executive Networks in Normal Aging and Mild Cognitive Impairment. Brain Connectivity, 7(7), 401–412. 10.1089/brain.2017.0509

10. Chang, C., Leopold, D. A., Schölvinck, M. L., Mandelkow, H., Picchioni, D., Liu, X., Ye, F. Q., Turchi, J. N., & Duyn, J. H. (2016). Tracking brain arousal fluctuations with fMRI. Proceedings of the National Academy of Sciences of the United States of America, 113(16), 4518–4523. 10.1073/pnas.1520613113

11. Chang, C., Liu, Z., Chen, M. C., Liu, X., & Duyn, J. H. (2013). EEG correlates of time-varying BOLD functional connectivity. NeuroImage, 72, 227–236. 10.1016/j.neuroimage.2013.01.049

12. Corbetta, M., & Shulman, G. L. (2002). Control of goal-directed and stimulus-driven attention in the brain. Nature Reviews. Neuroscience, 3(3), 201–215. 10.1038/nrn755

13. De Gennaro, L., Ferrara, M., & Bertini, M. (2001). EEG arousals in normal sleep: Variations induced by total and selective slow-wave sleep deprivation. Sleep, 24(6), 673–679. 10.1093/sleep/24.6.673

14. DuPre, E., Salo, T., Markello, R., Kundu, P., Whitaker, K., & Handwerker, D. (2020). ME-ICA/tedana: 0.0.9a [Computer software]. Zenodo. 10.5281/zenodo.3786890

15. Estarellas, M., Huntley, J., & Bor, D. (2024). Neural markers of reduced arousal and consciousness in mild cognitive impairment. International Journal of Geriatric Psychiatry, 39(6), e6112. 10.1002/gps.6112

16. Foucher, J. R., Otzenberger, H., & Gounot, D. (2004). Where arousal meets attention: A simultaneous fMRI and EEG recording study. NeuroImage, 22(2), 688–697. 10.1016/j.neuroimage.2004.01.048

17. Fransson, P. (2005). Spontaneous low-frequency BOLD signal fluctuations: An fMRI investigation of the resting-state default mode of brain function hypothesis. Human Brain Mapping, 26(1), 15–29. 10.1002/hbm.20113

18. Glasser, M. F., Coalson, T. S., Bijsterbosch, J. D., Harrison, S. J., Harms, M. P., Anticevic, A., Van Essen, D. C., & Smith, S. M. (2018). Using temporal ICA to selectively remove global noise while preserving global signal in functional MRI data. NeuroImage, 181, 692–717. 10.1016/j.neuroimage.2018.04.076

19. Glasser, M. F., Sotiropoulos, S. N., Wilson, J. A., Coalson, T. S., Fischl, B., Andersson, J. L., Xu, J., Jbabdi, S., Webster, M., Polimeni, J. R., Van Essen, D. C., & Jenkinson, M. (2013). The minimal preprocessing pipelines for the Human Connectome Project. NeuroImage, 80, 105–124. 10.1016/j.neuroimage.2013.04.127

20. Gonzalez, R., Gasco, J., & Llopis, J. (2019). University students and online social networks: Effects and typology. Journal of Business Research, 101, 707–714. 10.1016/j.jbusres.2019.01.011

21. Gonzalez-Castillo, J., Fernandez, I. S., Handwerker, D. A., & Bandettini, P. A. (2022). Ultra-slow fMRI fluctuations in the fourth ventricle as a marker of drowsiness. NeuroImage, 259, 119424. 10.1016/j.neuroimage.2022.119424

22. Greicius, M. D., Krasnow, B., Reiss, A. L., & Menon, V. (2003). Functional connectivity in the resting brain: A network analysis of the default mode hypothesis. Proceedings of the National Academy of Sciences, 100(1), 253–258. 10.1073/pnas.0135058100

23. Grimm, C., Duss, S. N., Privitera, M., Munn, B. R., Karalis, N., Frässle, S., Wilhelm, M., Patriarchi, T., Razansky, D., Wenderoth, N., Shine, J. M., Bohacek, J., & Zerbi, V. (2024). Tonic and burst-like locus coeruleus stimulation distinctly shift network activity across the cortical hierarchy. Nature Neuroscience, 27(11), 2167–2177. 10.1038/s41593-024-01755-8

24. Hansen, A. L., Johnsen, B. H., & Thayer, J. F. (2003). Vagal influence on working memory and attention. International Journal of Psychophysiology: Official Journal of the International Organization of Psychophysiology, 48(3), 263–274. 10.1016/s0167-8760(03)00073-4

25. Heitz, R. P., Schrock, J. C., Payne, T. W., & Engle, R. W. (2008). Effects of incentive on working memory capacity: Behavioral and pupillometric data. Psychophysiology, 45(1), 119–129. 10.1111/j.1469-8986.2007.00605.x

26. Huang, J., Sander, C., Jawinski, P., Ulke, C., Spada, J., Hegerl, U., & Hensch, T. (2015). Test-retest reliability of brain arousal regulation as assessed with VIGALL 2.0. Neuropsychiatric Electrophysiology, 1(1), 13. 10.1186/s40810-015-0013-9

27. Jacobs, H. I. L., Wiese, S., van de Ven, V., Gronenschild, E. H. B. M., Verhey, F. R. J., & Matthews, P. M. (2015). Relevance of parahippocampal-locus coeruleus connectivity to memory in early dementia. Neurobiology of Aging, 36(2), 618–626. 10.1016/j.neurobiolaging.2014.10.041

28. Jenkinson, M., Bannister, P., Brady, M., & Smith, S. (2002). Improved optimization for the robust and accurate linear registration and motion correction of brain images. NeuroImage, 17(2), 825–841. 10.1016/s1053-8119(02)91132-8

29. Jenkinson, M., Beckmann, C. F., Behrens, T. E. J., Woolrich, M. W., & Smith, S. M. (2012). FSL. NeuroImage, 62(2), 782–790. 10.1016/j.neuroimage.2011.09.015 Jones, B. E. (2003). Arousal systems. Frontiers in Bioscience: A Journal and Virtual Library, 8, s438–451. https://doi.org/10.2741/1074

30. Kim, A. J., Nguyen, K., & Mather, M. (2024). Eye movements reveal age differences in how arousal modulates saliency priority but not attention processing speed (p. 2024.05.06.592619). bioRxiv. 10.1101/2024.05.06.592619

31. Kramer, L., Sander, C., Bertsch, K., Gescher, D. M., Cackowski, S., Hegerl, U., & Herpertz, S. C. (2019). EEG-vigilance regulation in Borderline Personality Disorder. International Journal of Psychophysiology: Official Journal of the International Organization of Psychophysiology, 139, 10–17. 10.1016/j.ijpsycho.2019.02.007

32. Kundu, P., Brenowitz, N. D., Voon, V., Worbe, Y., Vértes, P. E., Inati, S. J., Saad, Z. S., Bandettini, P. A., & Bullmore, E. T. (2013). Integrated strategy for improving functional connectivity mapping using multiecho fMRI. Proceedings of the National Academy of Sciences, 110(40), 16187–16192. 10.1073/pnas.1301725110

33. Kundu, P., Inati, S. J., Evans, J. W., Luh, W.-M., & Bandettini, P. A. (2012). Differentiating BOLD and non-BOLD signals in fMRI time series using multi-echo EPI. NeuroImage, 60(3), 1759–1770. 10.1016/j.neuroimage.2011.12.028

34. Lendner, J. D., Helfrich, R. F., Mander, B. A., Romundstad, L., Lin, J. J., Walker, M. P., Larsson, P. G., & Knight, R. T. (2020). An electrophysiological marker of arousal level in humans. eLife, 9, e55092. 10.7554/eLife.55092

35. Liu, T. T., & Falahpour, M. (2020). Vigilance Effects in Resting-State fMRI. Frontiers in Neuroscience, 14. 10.3389/fnins.2020.00321 Marzano, C., Ferrara, M., Moroni, F., & De Gennaro, L. (2011). Electroencephalographic sleep inertia of the awakening brain. Neuroscience, 176, 308–317. https://doi.org/10.1016/j.neuroscience.2010.12.014

36. Mathersul, D., McDonald, S., & Rushby, J. A. (2013). Autonomic arousal explains social cognitive abilities in high-functioning adults with autism spectrum disorder. International Journal of Psychophysiology: Official Journal of the International Organization of Psychophysiology, 89(3), 475–482. 10.1016/j.ijpsycho.2013.04.014

37. Menon, V. (2011). Large-scale brain networks and psychopathology: A unifying triple network model. Trends in Cognitive Sciences, 15(10), 483–506. 10.1016/j.tics.2011.08.003

38. Menon, V., & Uddin, L. Q. (2010). Saliency, switching, attention and control: A network model of insula function. Brain Structure & Function, 214(5–6), 655–667. 10.1007/s00429-010-0262-0

39. Moehlman, T. M., de Zwart, J. A., Chappel-Farley, M. G., Liu, X., McClain, I. B., Chang, C., Mandelkow, H., Özbay, P. S., Johnson, N. L., Bieber, R. E., Fernandez, K. A., King, K. A., Zalewski, C. K., Brewer, C. C., van Gelderen, P., Duyn, J. H., & Picchioni, D. (2019). All-night functional magnetic resonance imaging sleep studies. Journal of Neuroscience Methods, 316, 83–98. 10.1016/j.jneumeth.2018.09.019

40. Mucha, P. J., Richardson, T., Macon, K., Porter, M. A., & Onnela, J.-P. (2010). Community Structure in Time-Dependent, Multiscale, and Multiplex Networks. Science, 328(5980), 876–878. 10.1126/science.1184819

41. Neal, J., Song, I., Katz, B., & Lee, T.-H. (2023). Association of intrinsic functional connectivity between the locus coeruleus and salience network with attentional ability. Journal of Cognitive Neuroscience, 35(10), 1557–1569. 10.1162/jocn_a_02036

42. Olbrich, S., Olbrich, H., Jahn, I., Sander, C., Adamaszek, M., Hegerl, U., Reque, F., & Stengler, K. (2013). EEG-vigilance regulation during the resting state in obsessive-compulsive disorder. Clinical Neurophysiology: Official Journal of the International Federation of Clinical Neurophysiology, 124(3), 497–502. 10.1016/j.clinph.2012.08.018

43. Oyarzabal, E. A., Hsu, L.-M., Das, M., Chao, T.-H. H., Zhou, J., Song, S., Zhang, W., Smith, K. G., Sciolino, N. R., Evsyukova, I. Y., Yuan, H., Lee, S.-H., Cui, G., Jensen, P., & Shih, Y.-Y. I. (2022). Chemogenetic stimulation of tonic locus coeruleus activity strengthens the default mode network. Science Advances, 8(17), eabm9898. 10.1126/sciadv.abm9898

44. Pedersen, M., Zalesky, A., Omidvarnia, A., & Jackson, G. D. (2018). Multilayer network switching rate predicts brain performance. Proceedings of the National Academy of Sciences, 115(52), 13376–13381. 10.1073/pnas.1814785115

45. Robison, M. K., & Brewer, G. A. (2020). Individual differences in working memory capacity and the regulation of arousal. Attention, Perception, & Psychophysics, 82(7), 3273–3290. 10.3758/s13414-020-02077-0

46. Robison, M. K., Ralph, K. J., Gondoli, D. M., Torres, A., Campbell, S., Brewer, G. A., & Gibson, B. S. (2023). Testing locus coeruleus-norepinephrine accounts of working memory, attention control, and fluid intelligence. Cognitive, Affective & Behavioral Neuroscience, 23(4), 1014–1058. 10.3758/s13415-023-01096-2

47. Rolls, E. T., Huang, C.-C., Lin, C.-P., Feng, J., & Joliot, M. (2020). Automated anatomical labelling atlas 3. NeuroImage, 206, 116189. 10.1016/j.neuroimage.2019.116189

48. Rubinov, M. (2016). Constraints and spandrels of interareal connectomes. Nature Communications, 7(1), 13812. 10.1038/ncomms13812

49. Rubinov, M. (2023). Circular and unified analysis in network neuroscience. eLife, 12, e79559. 10.7554/eLife.79559

50. Rubinov, M. (2025). Unifying equivalences across unsupervised learning, network science, and imaging/network neuroscience (No. arXiv:2508.10045). arXiv. 10.48550/arXiv.2508.10045

51. Sanda, P., Hlinka, J., van den Berg, M., Skoch, A., Bazhenov, M., Keliris, G. A., & Krishnan, G. P. (2024). Cholinergic modulation supports dynamic switching of resting state networks through selective DMN suppression. PLoS Computational Biology, 20(6), e1012099. 10.1371/journal.pcbi.1012099

52. Shine, J. M., Bissett, P. G., Bell, P. T., Koyejo, O., Balsters, J. H., Gorgolewski, K. J., Moodie, C. A., & Poldrack, R. A. (2016a). The Dynamics of Functional Brain Networks: Integrated Network States during Cognitive Task Performance. Neuron, 92(2), 544–554. 10.1016/j.neuron.2016.09.018

53. Shine, J. M., Koyejo, O., & Poldrack, R. A. (2016). Temporal metastates are associated with differential patterns of time-resolved connectivity, network topology, and attention. Proceedings of the National Academy of Sciences of the United States of America, 113(35), 9888–9891. 10.1073/pnas.1604898113

54. Shirer, W. R., Ryali, S., Rykhlevskaia, E., Menon, V., & Greicius, M. D. (2012). Decoding Subject-Driven Cognitive States with Whole-Brain Connectivity Patterns. Cerebral Cortex (New York, NY), 22(1), 158–165. 10.1093/cercor/bhr099

55. Smith, R., Keramatian, K., & Christoff, K. (2007). Localizing the rostrolateral prefrontal cortex at the individual level. NeuroImage, 36(4), 1387–1396. 10.1016/j.neuroimage.2007.04.032

56. Suttkus, S., Schumann, A., de la Cruz, F., & Bär, K. (2021). Working memory in schizophrenia: The role of the locus coeruleus and its relation to functional brain networks. Brain and Behavior, 11(5), e02130. 10.1002/brb3.2130

57. Teles-Grilo Ruivo, L. M., Baker, K. L., Conway, M. W., Kinsley, P. J., Gilmour, G., Phillips, K. G., Isaac, J. T. R., Lowry, J. P., & Mellor, J. R. (2017). Coordinated Acetylcholine Release in Prefrontal Cortex and Hippocampus Is Associated with Arousal and Reward on Distinct Timescales. Cell Reports, 18(4), 905–917. 10.1016/j.celrep.2016.12.085

58. Tsuno, N., Shigeta, M., Hyoki, K., Kinoshita, T., Ushijima, S., Faber, P. L., & Lehmann, D. (2002). Spatial organization of EEG activity from alertness to sleep stage 2 in old and younger subjects. Journal of Sleep Research, 11(1), 43–51. 10.1046/j.1365-2869.2002.00288.x

59. Uddin, L. Q. (2015). Salience processing and insular cortical function and dysfunction. Nature Reviews Neuroscience, 16(1), 55–61. 10.1038/nrn3857

60. Uddin, L. Q. (2016). Salience Network of the Human Brain. Academic Press.

61. Ulke, C., Huang, J., Schwabedal, J. T. C., Surova, G., Mergl, R., & Hensch, T. (2017). Coupling and dynamics of cortical and autonomic signals are linked to central inhibition during the wake-sleep transition. Scientific Reports, 7(1), 11804. 10.1038/s41598-017-09513-6

62. Ulke, C., Tenke, C. E., Kayser, J., Sander, C., Böttger, D., Wong, L. Y. X., Alvarenga, J. E., Fava, M., McGrath, P. J., Deldin, P. J., Mcinnis, M. G., Trivedi, M. H., Weissman, M. M., Pizzagalli, D. A., Hegerl, U., & Bruder, G. E. (2019). Resting EEG Measures of Brain Arousal in a Multisite Study of Major Depression. Clinical EEG and Neuroscience, 50(1), 3–12. 10.1177/1550059418795578

63. Unsworth, N., & Robison, M. K. (2017). A locus coeruleus-norepinephrine account of individual differences in working memory capacity and attention control. Psychonomic Bulletin & Review, 24(4), 1282–1311. 10.3758/s13423-016-1220-5

64. van der Wel, P., & van Steenbergen, H. (2018). Pupil dilation as an index of effort in cognitive control tasks: A review. Psychonomic Bulletin & Review, 25(6), 2005–2015. 10.3758/s13423-018-1432-y

65. Váša, F., & Mišić, B. (2022). Null models in network neuroscience. Nature Reviews. Neuroscience, 23(8), 493–504. 10.1038/s41583-022-00601-9 Xie, M., Huang, Y., Cai, W., Zhang, B., Huang, H., Li, Q., Qin, P., & Han, J. (2024). Neurobiological Underpinnings of Hyperarousal in Depression: A Comprehensive Review. Brain Sciences, 14(1), 50. 10.3390/brainsci14010050

66. Yerkes, R. M., & Dodson, J. D. (1908). The relation of strength of stimulus to rapidity of habit-formation. Journal of Comparative Neurology and Psychology, 18(5), 459–482. 10.1002/cne.920180503

67. Young, C. B., Raz, G., Everaerd, D., Beckmann, C. F., Tendolkar, I., Hendler, T., Fernández, G., & Hermans, E. J. (2017). Dynamic Shifts in Large-Scale Brain Network Balance As a Function of Arousal. The Journal of Neuroscience: The Official Journal of the Society for Neuroscience, 37(2), 281–290. 10.1523/JNEUROSCI.1759-16.2016

68. Zhang, S., Hu, S., Chao, H. H., Luo, X., Farr, O. M., & Li, C. R. (2012). Cerebral correlates of skin conductance responses in a cognitive task. NeuroImage, 62(3), 1489–1498. 10.1016/j.neuroimage.2012.05.036

